# Representational Geometries Across Visual Working Memory Encoding and Maintenance

**DOI:** 10.1101/2025.09.07.674590

**Authors:** Tomoya Nakamura, Seng Bum Michael Yoo, Kendrick Kay, Hakwan Lau, Ali Moharramipour

## Abstract

This study examined the representational geometry of visual working memory using human 7T fMRI. Observers viewed a face-scene blended sample, attended to either the face or scene, and judged whether the attended aspect matched a subsequent test image. We found that the coding dimension distinguishing faces and scenes rotated between encoding and maintenance phases, in both visual and association areas. In these regions, exclusive combinations (i.e., encoded face and maintained scene vs. encoded scene and maintained face) were linearly decodable, indicating encoding and maintenance phases represent visual information using distinct subspaces. Such high-dimensional, flexible geometry can, in principle, protect maintained visual information from incoming input. At the same time, robust cross-decoding was observed across phases, reflecting the stability and generalizability of the represented contents. The balance between representational flexibility and stability varied along the cortical hierarchy: early sensory regions emphasized flexibility, whereas transmodal regions showed greater stability.

## Introduction

How working memory maintains information has long been a central question in cognitive neuroscience. The classical view, largely derived from monkey electrophysiology, posits that working memory content is supported by persistent neural activity in the prefrontal cortex that maintains a stable code over time throughout the delay period (e.g., [1,2]). Early human fMRI studies (e.g., [3–5]) primarily relied on univariate analyses of the delay-period activity to explore neural correlates of working memory under the assumption that sustained activity during this period reflects maintained information.

More recent work, however, has challenged this static view, proposing instead that working memory representations evolve over time, not simply in terms of activity magnitude, but also more fundamentally in the way information is coded, which may help minimize interference from ongoing sensory input (e.g., [6–11]). In animal electrophysiological studies, these transformations have been conceptualized in terms of rotational dynamics, in which neural representations of task-relevant information rotate along consistent, structured trajectories in the underlying neural population state space (e.g., [12–14]).

Such perspectives motivate analyzing working memory through representational geometry, where neural population activity is characterized by the relative arrangement of activity patterns in high-dimensional space (e.g., [15–19]).

Specifically, working memory representations can be characterized as changes in the orientation and dimensionality of the subspace formed by neural population activity patterns (Figure 1). A key question, therefore, is how such transformations preserve the integrity of the information across working memory phases while altering the underlying representational format.

**Figure 1.**
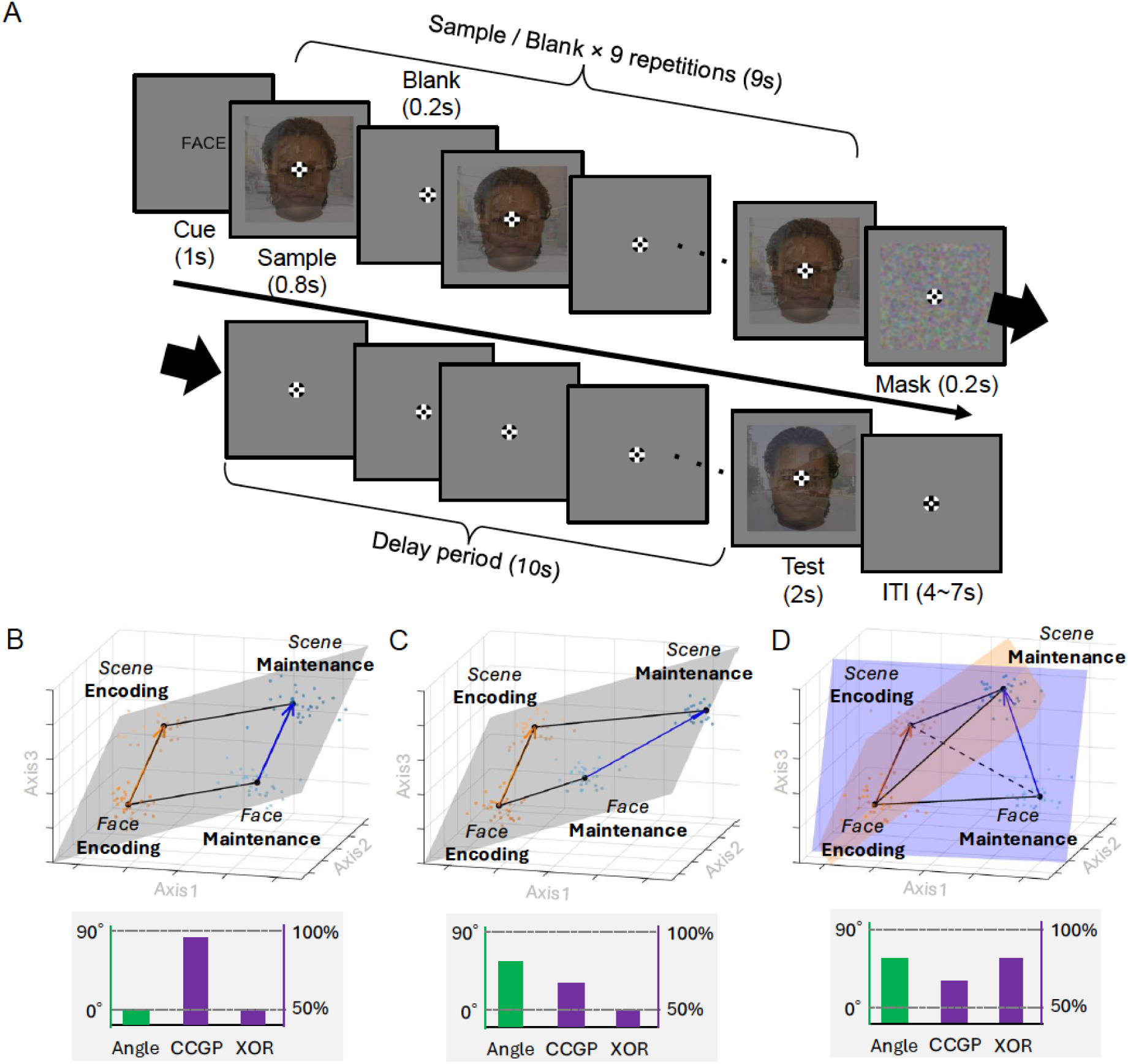
Task design and analytic framework of representational geometry. (A) In each trial, observers were instructed to memorize only the cued aspect (either the face or the scene) of a face-scene blended sample stimulus, which was repeatedly presented (encoding phase). Following a delay period (maintenance phase), a test stimulus appeared in a randomly selected half of the trials, and observers judged whether the cued aspect of the test stimulus was the same as or different from the sample. In the remaining half of the trials, no test stimulus was presented, and no response was required. In this example of a face-cued trial, the face remains the same while the scene changes; therefore, the correct response is "SAME." Note that because only the cued aspect (face vs. scene) is manipulated, the paradigm can more effectively reflect encoding-related processing rather than raw stimulus-driven activity. The face and scene stimuli are reproduced with permission from Ma et al. [32] and Konkle et al. [33]. Abbreviations: ITI, intertrial interval. (B) A hypothetical representational space where the coding axes distinguishing faces from scenes are parallel across encoding (orange arrow) and maintenance phases (blue arrow). In this case, the angle between the coding axes is zero, leading to perfect generalization as indicated by cross-condition generalization performance (CCGP). However, linear, binary exclusive-or (XOR) decoding (i.e. binary classification of encoded face and maintained scene vs encoded scene and maintained face) would be impossible. (C) A case in which the coding axis rotates within the same subspace over time. All four points corresponding to each condition still lie on the same plane (gray). In this case, the angle between coding axes is non-zero, CCGP decreases, and XOR decoding remains impossible. (D) A case where the coding axis rotates to define distinct subspaces for encoding (orange plane) and maintenance phases (blue plane). In this case, the angle between coding axes is non-zero, CCGP decreases, and XOR decoding is above chance. Note that here the axes represent three example effective axes within a higher-dimensional space.

Importantly, working memory representations are not confined to the prefrontal cortex. Accumulating evidence, especially from multivariate decoding, indicates that working memory content is distributed across a broad network of brain regions, including occipitotemporal cortex, posterior parietal cortex, and lateral prefrontal cortex (e.g., [20–25]; for a review, see [26–28]). In contrast to animal neurophysiology, a large body of human fMRI work was focused on multivoxel patterns in the visual cortex, often using cross-decoding approaches to compare encoding- and maintenance-stage representations. For example, Harrison and Tong [29] demonstrated that stimulus orientations maintained in working memory could not only be decoded from the early visual cortex, but also successfully crossdecoded from stimulusevoked activity within the same regions (see also [30,31]). These findings indicate that sensory regions retain shared visual representation between perceptual and mnemonic states.

Taken together, it is possible that different brain regions exhibit distinct transformation profiles across the encoding and maintenance phases. For example, some regions may show more generalizable representations across phases, whereas others may exhibit greater transformations forming distinct subspaces. However, the largescale organization of such representational geometry across the entire cortex remains unexplored.

In the present study, we used 7T fMRI to examine how the representational geometry of visual stimuli evolves across working memory phases while participants encoded and maintained either faces or scenes. Extending prior region-specific approaches (such as in most animal neurophysiological studies), we characterized these transformations across the whole cortex to reveal organization principles of working memory representations. To anticipate, we found that representations during encoding and maintenance occupy non-parallel subspaces, expanding dimensionality, while still preserving partial cross-phase generalization. This balance tended to vary along the cortical hierarchy, with unimodal regions exhibiting greater transformation than transmodal regions.

## Results

### Working memory representations rotated across encoding and maintenance phases

Using ultra-high-field (7T) fMRI, we measured cortical activity while observers (n = 8) performed a delayed-match-to-sample task on face–scene blended stimuli (Figure 1A). On every trial, a verbal cue ("FACE" or "SCENE") instructed which aspect of the upcoming sample to attend and remember. A face–scene blended image was then presented (encoding phase), followed by a delay (maintenance phase). On a randomly selected half of the trials, a test image was presented at the end and observers were asked to judge whether the cued aspect had changed between the sample and the test. Behavioral performance was high (mean correct response rate = 94.3%), indicating that observers followed the instructions and were well engaged in the task (Figure S1).

One key feature of our design is that both facecued and scenecued trials showed the *same* blended sample and differed only in which aspect was cued. In this way, the encoding phase reflects attention-driven representation rather than a lowerlevel stimulusdriven sensory activity. It is also important to note that all of our subsequent analyses operate on the face−scene contrast within each phase, and on how the resulting coding axis alters across phases. We therefore focus on the interaction between the working memory process (encoding vs. maintenance) and visual category (face vs. scene), rather than the main effect of either factor. In this format, superficial sensory differences are effectively canceled out by the facescene contrast within each phase, allowing the analysis to focus on differences in representational formats across encoding and maintenance phases. Consequently, the design’s apparent asymmetry (i.e., the presence of the blended image, as well as its unattended aspect during the encoding phase and no image during the maintenance phase), would not, by itself, bias the findings. Overall, the use of blended images helps better focus on encoding-related processing (rather than on raw stimulus-driven activity), and the analysis approach ensures valid comparisons with the maintenance phase.

We extracted vertex-level wholebrain BOLD activity from 22 brain regions-of-interest (ROIs) defined by the Glasser atlas [34] under four conditions: encoded face, encoded scene, maintained face, and maintained scene (Figure S2). These corresponded to working memory activity when observers selectively encoded the face or scene aspect of the sample and when they subsequently maintained the cued aspect during the delay. In a neural state space in which each vertex serves as a dimension, we defined two separate face–scene coding axes, one for the encoding and one for the maintenance phase, as the lines connecting the multivertex beta patterns for face- and scene-cued trials (scene beta pattern − face beta pattern) (Figure 1B).

We first evaluated the robustness of face-vs-scene multivertex activation pattern within each ROI by applying a testretest reliability approach (e.g., [21]). For each encoding and maintenance phase, split-half correlations were computed on face–scene difference patterns. Despite substantial individual variability, we confirmed that nearly all ROIs showed significant reliability for both phases at the group level (Figure S3).

On this basis, we quantified the geometric relationship between representations during encoding and maintenance phases by measuring the angle between the face-vs-scene coding axes across phases. When the two axes are parallel (angle = 0°), both phases use the same multivariate dimension to distinguish faces from scenes (Figure 1B); when they are orthogonal (90°), they use independent dimensions; intermediate angles indicate that the coding scheme has partially rotated over time (Figure 1C, D).

Because finite-sample noise causes even truly parallel axes to deviate from 0°, with the magnitude depending on each ROI’s signal-to-noise ratio (SNR), we compared the observed angle against a noise floor: the angle expected between two parallel axes under SNR matched to that of the observed axes (Figure S4).

We calculated the z-values of the angle deviation from the noise floor to perform a formal statistical test. All but three ROIs (somatosensory/motor, paracentral/mid cingulate, and posterior opercular) exhibited significant rotation at the group-level, indicating that rotation over different phases in working memory processing is prevalent among distributed brain regions, irrespective of processing hierarchy (Figure 2A).

**Figure 2.**
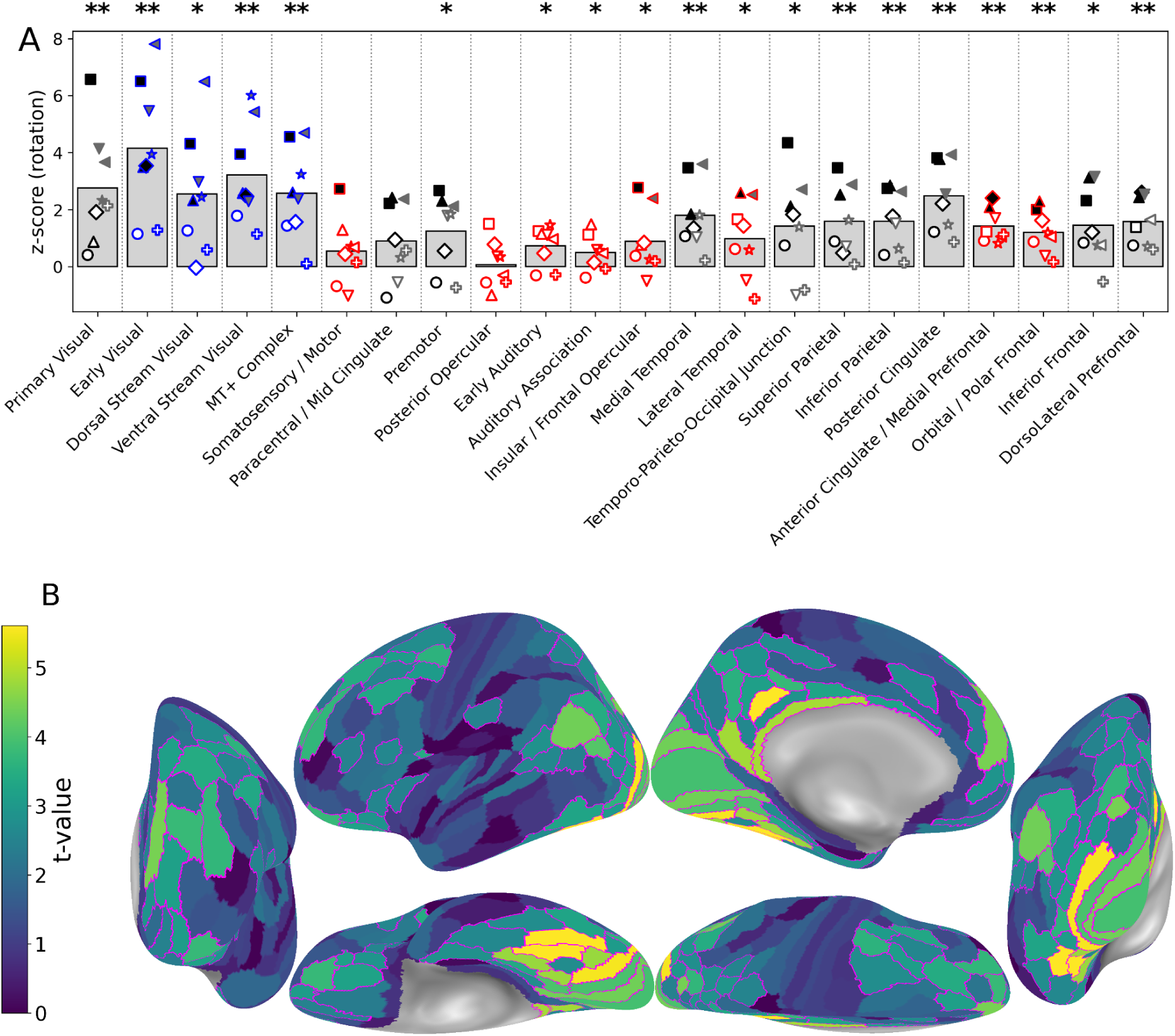
Rotation of the coding axis between encoding and maintenance phases. (A) z-values reflecting deviation from the noise floor (parallelism under noise; Figure S4) are shown for all observers and ROIs. Bar graphs indicate the inter-individual mean. Filled markers indicate observers who exhibit individual-level significance (FDR-corrected), whereas unfilled markers indicate those who do not. Black and gray markers indicate pilot and new observers, respectively (see the Methods section). ROIs marked with blue and red indicate universality and sporadicity of significant individuals, respectively. Asterisks denote FDR-corrected group-level significance (**q < .01, *q < .05). At the group-level, the coding axis rotation was evident in the visual and some parietal/frontal association areas (for detailed statistics, see Table S1). Note that other nonsignificant regions do not necessarily have parallel coding axes but simply noisy visual representation (see also Figure S4). (B) Fine-grained surface maps illustrating significant axis rotation. Group-level t-values are visualized on the Glasser atlas, with negative t-values clipped to zero owing to the one-tailed nature of the test. ROIs showing significant rotation (q < .05) are outlined with purple contours.

Even at the individual level, we found a reliable rotation, especially in the visual cortex. The universality test [35] revealed that the significance of rotation in most visual areas, including early visual, dorsal/ventral stream visual, and MT+, was universal across observers (i.e., reached individual-level significance in at least 5 of the 8 observers). In contrast, in addition to the three non-significant ROIs at the group-level, we found the significance of the rotation in early auditory, auditory association, insular/frontoopercular, lateral temporal, anterior cingulate/medial prefrontal, and orbital/polar frontal areas was sporadic (i.e., at least 6 observers did not exhibit individual-level significance). The regions exhibiting reliable rotation were also consistent when tested using fine-grained parcellations (Figure 2B). Note that the lack or sporadicity of significance does not necessarily imply parallel axes; rather, visual tuning in these regions was too noisy, thereby elevating the noise floor of the angle (Figure S4).

### Encoding and maintenance representations occupied different subspaces

Theoretically, the rotation could occur within the same subspace (Figure 1C), or it could involve distinct subspaces, leading to an increase in the effective dimensionality of the representational space (Figure 1D). In order to distinguish these alternative scenarios, we evaluated the dimensionality by assessing the linear separability of variables. When multiple variables are represented, the greater the number of dichotomies that can be separated by linear hyperplanes, the higher the effective dimensionality [36]. Specifically, when rotation of a coding axis recruits an extra, new dimension (as described in Figure 1D), this expansion can facilitate flexible coding and thereby enable decoding of an exclusive-or (XOR) dichotomy [15]. In our case, where two factorial variables each have two levels (e.g., the combination of content: *face* vs. *scene* and processing phases: *encoding* vs. *maintenance*), XOR contrasts Class A (*encoded face* together with *maintained scene*) against Class B (*encoded scene* together with *maintained face*).

To test for XOR decodability, we obtained two beta vectors from each trial: one from the encoding phase and one from the maintenance phase, so every trial, depending on the cue, supplied one beta to Class A and one to Class B. A logistic-regression classifier was then trained to discriminate the two classes, with training and test data drawn from independent runs (see Methods). Above-chance XOR decoding requires the four cue × phase combinations to be arranged in a three-dimensional geometry (Figure 1D).

We found that the XOR dichotomy was decodable above-chance from all ROIs except the auditory regions (Figure 2). Regions with strong XOR decodability largely overlapped with those showing substantial rotation of the face-vs-scene coding axes (Figure 2). This pattern indicates that the face-vs-scene coding axes do not merely rotate within a single representational space, but instead define at least partially distinct subspaces during the maintenance phase.

### Working memory representations were generalizable across encoding and maintenance phases

Although rotation of the coding axis expanded the dimensionality, it was still possible that two coding schemes are not fully orthogonal and therefore remain generalizable to each other. To test this possibility, we used Cross-Condition Generalization Performance (CCGP) [15] to quantify how well the linear decoder trained in one phase could generalize to the other phases (from encoding to maintenance and vice versa). Higher CCGP values reflect coding that is more invariant across phases.

First, face vs. scene was decodable from all ROIs except the early auditory cortex (Figure 3; even separately within encoding and maintenance phases, see Figure S5), indicating that visual content was represented across widespread cortical regions. Among these content-sensitive regions, we observed above-chance CCGP except for the primary visual cortex. This indicates that most areas coded the content information in a partially invariant (i.e., generalizable) format across encoding and maintenance phases (Figure 3).

**Figure 3.**
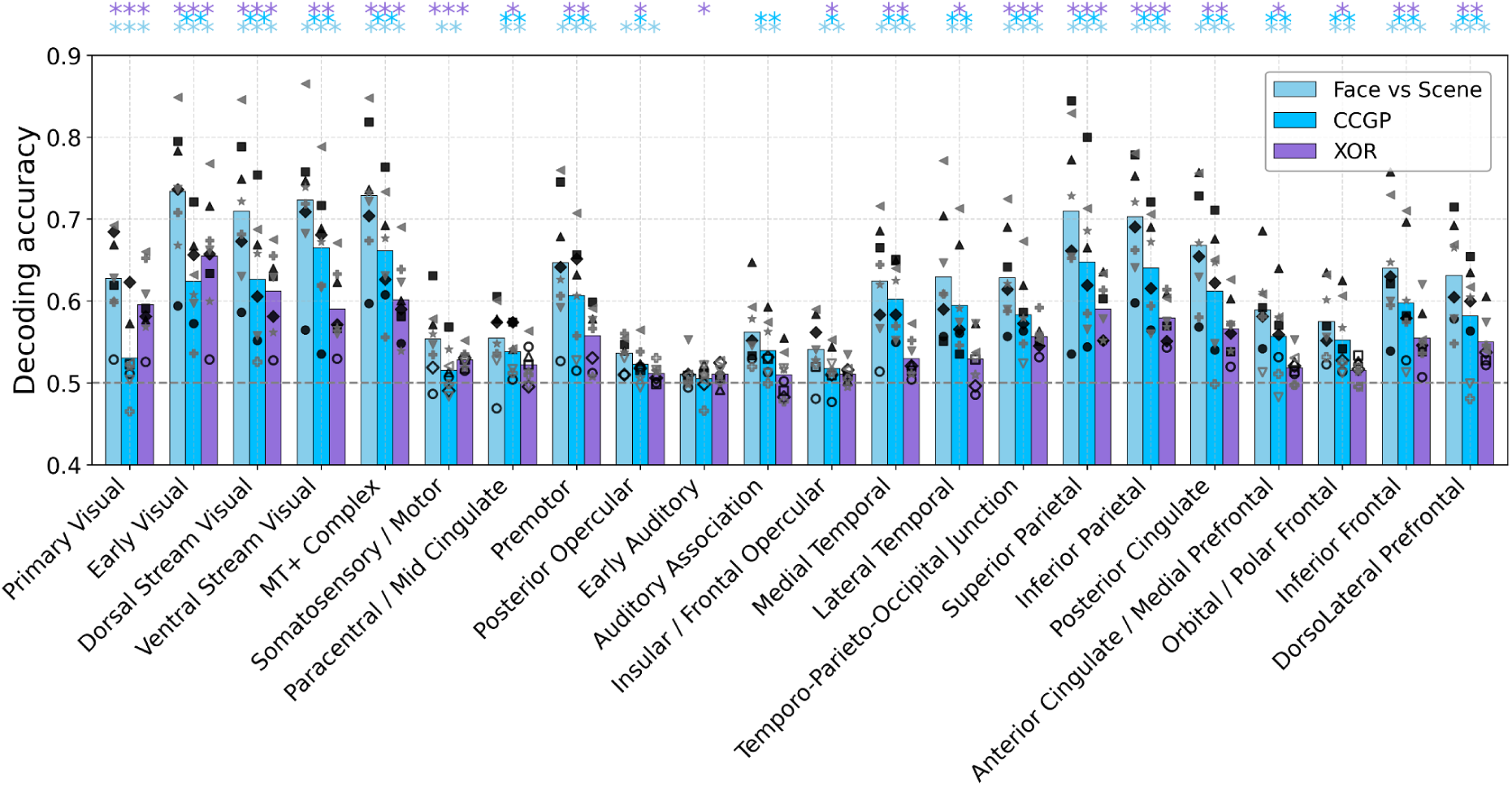
Face-vs-scene decoding, cross-condition generalization performance (CCGP) and exclusive-or (XOR) decoding. Decoding accuracy between face-cued and scene-cued trials, regardless of phases (i.e., encoding and maintenance), is shown in light blue, CCGP for face–scene dichotomy (trained to decode face vs. scene in one phase and tested in the other) is shown in deep blue, and XOR (decoding encoded face and maintained scene vs. encoded scene and maintained face) is shown in purple. Individual observer data are plotted around the bars (interindividual mean) using markers of different shapes. Filled markers indicate individual-level significance assessed by permutation tests. Black and gray markers indicate pilot and new observers, respectively (see the Methods section). Asterisks denote the significance of the group-level one-sample t-test (one-tailed) on z-transformed accuracy, FDR-corrected across all ROIs (**** q < .0001, *** q < .001, ** q < .01, * q < .05). The face-scene dichotomy was decodable except for the early auditory cortex and generalizable across phases in all regions except for the primary visual cortex, somatosensory/motor cortex, and early auditory cortex. Therefore, visual content is represented in a manner generalizable over different working memory processing phases in all areas except for early sensory-motor areas. XOR dichotomy was decodable from all visual areas and most association areas (e.g., temporal, parietal, and prefrontal regions), indicating partially different subspaces for encoded and maintained content (for detailed statistics, see Table S2, S3, and S4). When the runs with poor image quality were excluded, XOR and CCGP in the insular/frontal opercular cortex became non-significant. All decoding analyses were performed using training and test data from independent runs.

We should note that residual within-trial BOLD overlap between encoding and maintenance phases, if present, could potentially inflate the CCGP. Importantly, CCGP was computed with training and test data drawn from independent runs (see Methods), so above-chance cross-decoding cannot at least be attributed to within-run overlap of slow BOLD responses. In addition, collinearity between encoding- and maintenance-phase regressors remained within a controlled range (encoding phase: variance inflation factor (VIF) = 2.14; maintenance phase: VIF = 1.86), indicating that the two epochs were sufficiently dissociable at the design-matrix level.

Notably, such BOLD overlap would tend to make the two coding axes appear more parallel, not less, so the rotation and XOR decoding accuracy reported in the previous subsections are actually conservative with respect to this potential confound, making it unlikely to generate false positives. Taken together, although the hemodynamic contamination across phases cannot be fully accounted for, these considerations suggest that the present findings are unlikely to be strongly influenced by it.

### Representational formats were more stable in the association cortex compared to the early sensory regions

The ROIs with greater XOR decoding tended to exhibit larger cross-decoding performance drops (Figure 3). Such a tradeoff between flexible and stable geometry is theoretically predicted [15]. Importantly, here we provide evidence for the co-occurrence of representational flexibility and stability in human fMRI data, as XOR decodability did not eliminate CCGP in most ROIs.

Interestingly, regions located at the sensory–motor periphery of the cortical hierarchy (i.e., primary/early visual and somatomotor cortices) tended to prioritize flexible formats at the expense of stable formats (Figure 4A). In contrast, higher-tier visual regions (especially for the ventral visual stream) and association cortices (especially for the temporal cortex) showed the opposite pattern, exhibiting relatively greater stable geometry. This organizational trend was particularly evident along the visual pathways, when the data were visualized on a fine-grained atlas (Figure 4B).

**Figure 4.**
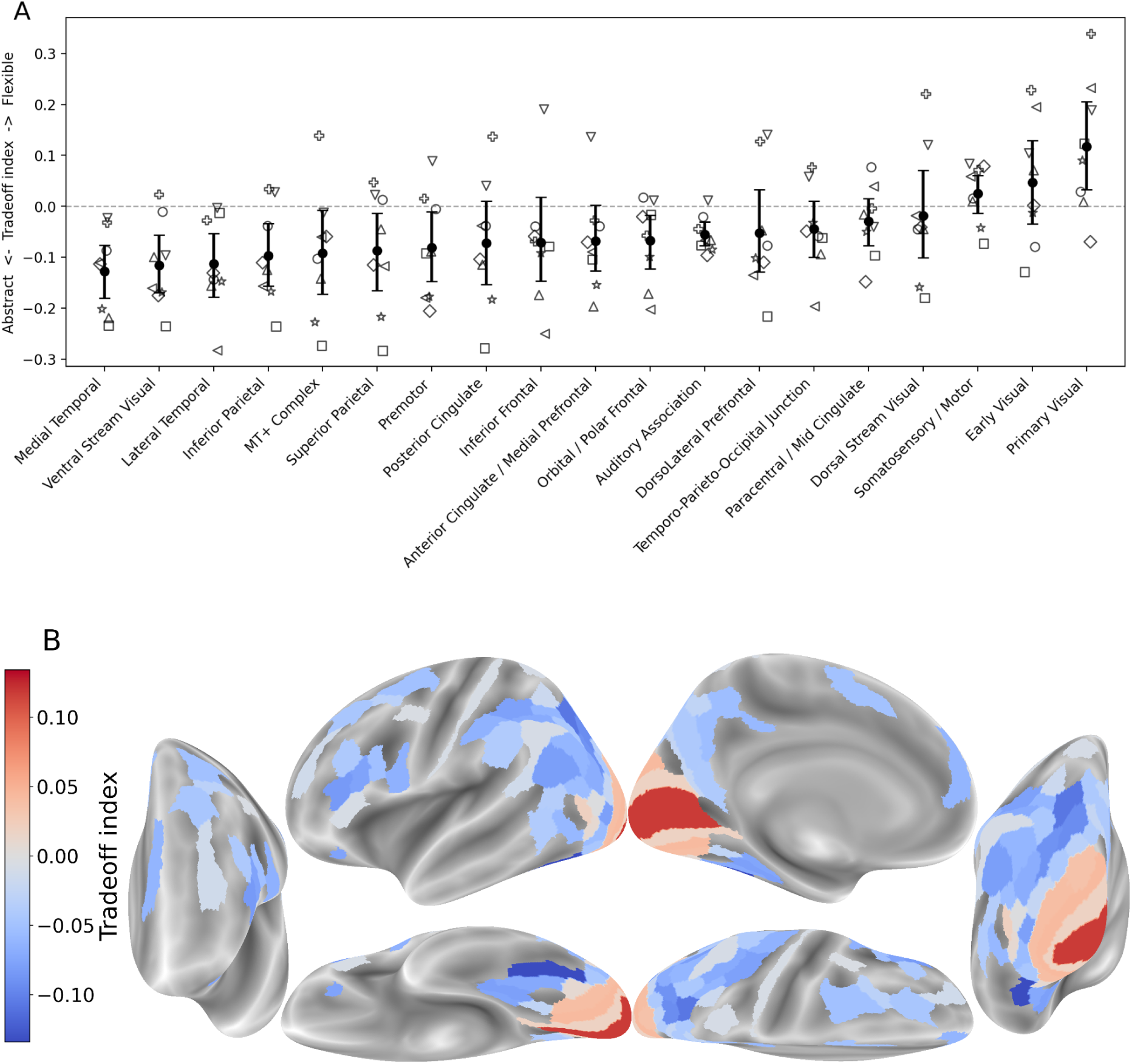
The level of tradeoff between stable and flexible geometry. (A) Tradeoff index for each of the 22 coarse-grained ROIs (shown in ascending order). Data show the inter-observer mean with 95% bootstrapped confidence intervals, with individual observers plotted as distinct marker shapes. A positive index indicates that XOR decoding exceeds CCGP, reflecting a tendency toward flexible formats at the cost of stable formats. (B) The tradeoff index derived for the fine-grained cortical atlas. ROIs with poor face-vs-scene decoding accuracy (q > .01) were excluded because they were not reliably tuned to the face–scene distinction. ROIs with poor CCGP and XOR performance (q > .01 for both measures) were also excluded because their tradeoff index could not be reliably estimated. Overall, unimodal regions (e.g., early visual areas) tended to favor flexible formats by separating encoding and maintenance subspaces, whereas transmodal regions exhibited a more balanced tradeoff or favored stable formats.

Notably, this trend appeared to align with the principal functional gradient from unimodal to transmodal cortex, previously identified in resting-state functional connectivity [37] and in multitask representational similarity [38], as shown in Figure S6; although, it did not survive statistical testing under a stringent null model that preserved spatial autocorrelation among parcels [38,39].

## Discussion

Using 7T fMRI across multiple sessions, we investigated how working memory representations evolve from encoding into maintenance phases. We found that, across multiple cortical regions, including the occipitotemporal, posterior parietal, and lateral prefrontal cortices, the representational geometry of task-relevant visual information was preserved across encoding and maintenance phases (as evidenced by cross-decoding performance; CCGP in Figure 3), while also occupying partially distinct subspaces (as evidenced by XOR decodability in Figure 3). These findings suggest that working memory does not simply preserve visual representations in a static form, but instead balances representational stability and transformation.

Although face–scene decoding was successful across cortical regions, this does not necessarily imply that all regions explicitly encoded face or scene identity *per se*.

Unlike canonical category-selective regions such as the fusiform face area (FFA) and parahippocampal place area (PPA) [40,41], some regions may instead distinguish the two conditions through other visual or cognitive features. The following aspects may help constrain what kind of information were likely contributed to face–scene decoding. First, because observers needed to detect unpredictable changes after the delay period, successful task performance likely required maintenance of the entire visual content rather than isolated low-level attributes alone. Second, scene subcategories (natural, manmade, open, and closed) could be decoded during the task (Figure S7), suggesting that the representations contained finer-grained content information beyond coarse attentional-state differences [42] or simple verbal labels of “FACE” versus “SCENE”.

Importantly, it is challenging to interpret differences in decoding performance across ROIs as they often differ in (raw and functional) SNR, microanatomy, functional topography, and neurovascular coupling (e.g., visual cortex vs. prefrontal cortex; see [43–45]). For instance, a lower cross-decoding is naturally expected in ROIs with lower baseline decoding performance for the reasons above. Therefore, to fairly quantify properties of the representational geometry across the entire cortex, we assessed CCGP (bidirectional cross-decoding) in conjunction with XOR decodability. Within the “geometry of abstraction” framework [15, 46–49], CCGP captures the extent to which neural representations generalize across conditions, whereas XOR decodability reflects high-dimensional coding that supports flexible separation of context-dependent information. We also measured the absolute rotation, which could capture transformation of representation formats even when it did not substantially increase dimensionality and therefore did not yield XOR decodability. By jointly assessing these complementary geometric measures, we were able to characterize regional differences and cortical organization more fairly in terms of the balance between representational flexibility and stability, rather than relying on simple differences in decoding or cross-decoding performance.

The early visual cortex showed particularly strong subspace rotation (substantially greater XOR than CCGP) between encoding and maintenance phases. Importantly, this finding does not preclude a contribution of the early visual cortex to working memory, as proposed by the sensory recruitment hypothesis ([31, 50–53], but see also [23,54]). Rather, it suggests that maintained representations in the sensory cortex are not merely degraded versions of initially encoded representations, but instead qualitatively different formats [53–60]. Such segregation may help protect mnemonic information from interference by ongoing sensory input [12,55]. One potential implementation is layer-specific separation: bottom-up input primarily targets the middle layers, whereas maintained content in working memory may rely more on top-down signals (see [61]), which are conveyed through deep and superficial layers [62–64].

Another possibility, which is not mutually exclusive with the above, is that the strong subspace rotation observed in the early visual cortex partly reflects differences in the spatial precision of perceptual and mnemonic representations. Because neurons in the early visual cortex possess small receptive fields, working memory representations with reduced spatial precision (e.g., [65,66], but see [67]) may recruit partially different neuronal populations from those engaged during sensory encoding. A similar principle may also extend to higher-level visual regions, where differences in spatial precision may explain the recruitment of distinct subpopulations of neurons across encoding and maintenance [68]. Future studies may help dissociate these possibilities by comparing the receptive field characteristics of vertices preferentially engaged during encoding and maintenance phases (e.g., [69]).

In contrast, association cortices showed greater generalization across encoding and maintenance phases despite still exhibiting partial subspace rotation. These findings suggest that association cortex favors more abstract and stable coding strategies.

Consistent with this, previous studies have shown that working memory representations in the parietal cortex remain relatively stable even in the presence of distractors [70]. Electrophysiological findings from the lateral prefrontal cortex [71] suggest this balance between format stability and geometric flexibility in association areas may arise from intermixed populations of neurons—some tuned selectively to perceptual or mnemonic content and others tuned to both (see also [72,73]).

Notably, the balance between representational flexibility and stability appeared to vary systematically along the unimodal–transmodal cortical hierarchy. Unimodal regions tended to exhibit stronger subspace rotation at the expense of stability, whereas geometry in transmodal regions was relatively more stable over time. This observation extends previous gradient-based accounts of cortical organization [36,37,74] by embedding them within a representational geometry framework. From this perspective, the unimodal–transmodal gradient may reflect a large-scale organizational principle governing how neural systems allocate representational resources between context-dependent and context-invariant coding. Greater stability in transmodal regions is also consistent with the accounts proposing progressively longer temporal windows along the cortical hierarchy, thereby supporting representations that keep and integrate information over extended timescales (e.g., [75,76]).

A key difference from the previous fMRI studies is that our goal was not to contrast perception and working memory as separate cognitive states, but rather to examine how working memory representations evolve from initial sensory encoding into subsequent maintenance. Accordingly, encoding and maintenance phases were intentionally embedded within the same trial to preserve the continuity of working memory processing, unlike prior studies that often used separate perception-only trials.

This design choice, however, introduces a potential limitation. Because encoding and maintenance always occurred in temporal succession, face-scene contrast during the maintenance-phase may have contained residual contributions from encoding-phase due to the sluggish nature of the hemodynamic response [28]. However, observed cross-phase generalization is unlikely to be strongly driven by the overlap of sluggish BOLD response, because collinearity of regressors for encoding and maintenance phases were modest (i.e., VIF was around 2). In addition, when computing CCGP, we used encoding and maintenance activity patterns derived from independent groups of runs as a safeguard against this potential issue. Although, we may not be able to completely rule out this fMRI-specific limitation, it is also important to note that encoding and maintenance may not constitute fully discrete neural states in the first place, as contemporary accounts increasingly emphasize continuous and dynamically evolving working memory representations [6–11]. On the other hand, such BOLD overlap would be expected to naturally reduce rather than inflating estimates of angle and XOR decodability. Therefore, the observed rotation and dimensionality expansion however cannot be attributed to such hemodynamic overlap. Moreover, even if the residual BOLD overlap biases representations toward greater apparent similarity across phases, it is unlikely to explain the relative differences in representational regime observed across cortical regions, which constitute one of our key findings (Figure 4).

Another notable difference from previous studies concerns the role of attention during encoding. Earlier studies often diverted observers’ attention to secondary tasks during perceptual conditions (e.g., letter identification at fixation; see [29]), raising the possibility that apparent differences between encoding- and maintenance-phase representations were partly influenced by attentional factors (for an exception, see [77]). In contrast, the present study defined the face-scene contrast solely by task-relevance, thereby more closely matching attentional states across working memory phases.

Moreover, because only a single isolated stimulus was typically presented in previous studies [20,29,30,71], decoding during the encoding phase may have been strongly driven by stimulus-driven sensory activity rather than by cognitive processes operating on the stimulus. In contrast, in the current study, by presenting overlaid face–scene stimuli and matching visual input across face-cued and scene-cued trials, the face–scene contrast was independent of stimulus-driven activity regardless of phases.

In a related line of research, Dijkstra et al. [78] investigated how the human brain distinguishes or confuses visual perception and visual imagery—a form of internally generated representation that is conceptually and neurally similar to visual working memory [29,79,80]. They showed that the behavior and neural responses in fusiform regions were best explained by a model in which congruent imagery and perception summate along a single dimension, such that excessively vivid imagery can be subjectively experienced as real even in the absence of sensory input.

By contrast, working memory representations are unlikely to be mistaken for real on their own. Rather, they are more plausibly expressed as contaminating or biasing representation of concurrently encoded perception of external stimuli [81,82], making it more necessary for the brain to keep them separable. Although it remains unclear which brain regions are fundamentally related to subjective experience, we consider that subspace rotation may be sufficient to reduce subjective confusion between contents that are concurrently being encoded and maintained ([49,83]; see also [84]).

At the same time, however, working memory phenomenology, if any, should remain continuous with that accompanying stimulus encoding, allowing internally maintained contents to be attributed to the same external source. From this perspective, some degree of representational generalization across encoding and maintenance phases may also be functionally important. One intriguing possibility is that individual differences in working-memory strategy, particularly the extent to which observers rely on vivid visual imagery, may modulate the balance between representational generalization and transformation. Observers relying more strongly on imagery-like strategies may maintain representations in formats that are more stable across phases.

It is noteworthy that one limitation of our study is that we did not directly establish the link between subspace rotation and behavior. Thus, it may be informative for future studies to titrate task difficulty (e.g., through parametric manipulation of synthesized naturalistic stimuli; see [60]) to yield a sufficient number of incorrect trials. This would perhaps enable trial-by-trial prediction of response accuracy based on the degree of subspace rotation in specific brain regions. Alternatively, future work could employ behavioral paradigms that introduce distractors during the delay period and also allow for higher-resolution reports, such as continuous reproduction tasks [81]. Such designs may further allow formalization of models of subjective experience—for example, how working memory representations are contaminated by concurrent perceptual input—based on the geometric relationship between working memory and perceptual representations.

## Methods

### Pre-registration

The eight observers analyzed in this study comprise four pilot observers, whose data were initially reported in a bioRxiv preprint [https://www.biorxiv.org/content/10.1101/2025.09.07.674590v1], and four additional observers recruited after the pre-registration of the analysis plan. The pre-registered plan was deposited as part of that preprint after the four pilot observers had been collected and analyzed.

The pre-registered components are: (i) the experimental protocol and stimulus parameters described below; (ii) the three measures of interest — coding-axis rotation, CCGP for the face–scene dichotomy, and XOR decoding; (iii) the stopping rule for data collection (see Observers); and (iv) the set of ROIs. Unless otherwise noted, all analyses and experimental procedures described below were conducted in accordance with the pre-registered plan.

### Observers

Eight naïve observers and one author (four males and five females; aged 21–28 years) participated in our task. We excluded one observer’s data from the analyses, as they were able to complete only one out of four scanning sessions. Except for that observer, all the other eight observers completed four scan sessions, on different days.

All observers reported that they have normal or corrected-to-normal visual acuity, normal color vision and no history of neuropsychiatric disorders. The study was approved by the Institutional Review Board at Sungkyunkwan University. We obtained written informed consent from all observers before the experiment.

As pre-registered, observer recruitment was terminated based on the universality of individual-level significant effects [35]. After collecting data from eight observers (the pre-registered minimum sample size), we met the pre-registered stopping criterion: for each of the three measures of interest (coding axis rotation, CCGP for face–scene dichotomy, and XOR decoding), more than half of the 22 ROIs reached the threshold for sufficient universality (likelihood ratio > 50 following [35]), regardless of whether the specific ROIs overlap across measures. Assuming a reasonable sensitivity of 1 – β = .9, this corresponded to finding at least 5 out of 8 observers showing significant effects or finding at least 6 out of 8 observers showing non-significant effects (with α = .05, FDR-corrected across ROIs).

### Apparatus

The experimental code was written in MATLAB R2021b (MathWorks, Natick, MA, USA) with the Psychphysics Toolbox 3.0.19 [85,86] and was run on a Ubuntu-based laptop. Visual stimuli were displayed on a screen behind the scanner and observed by observers through a mirror attached to the head coil. The refresh rate and spatial resolution of the monitor was 60 Hz and 1600 × 1000 pixels, respectively. The effective viewing distance was 104 cm.

### Stimuli

We presented identical sets of face–scene blended sample images and manipulated only the aspect (face vs. scene) that was attended and remembered. In this way, the contrast between face-cued and scene-cued trials—which is our main focus—reflects primarily the difference in perceptual or mnemonic representation under matched stimulus input.

To create the set of face-scene blended sample images, first we selected 32 different stimuli (4 sub-categories × 8 exemplars) each for faces and scenes. These numbers were chosen to balance the diversity of the stimuli with the number of repetitions of the same stimuli. Diversity was expected to help extract the general representation of face/scene and minimize the longer-term familiarity effect that can confound working memory. In contrast, repetition was expected to improve data reliability.

Face stimuli were drawn from the Chicago Face Database [32], comprising 4 sub-categories defined by the combination of race and sex (Black/Female, Black/Male, White/Female, and White/Male), with all faces displaying neutral expressions. Scene stimuli were drawn from the “Massive Memory” dataset [33], comprising 4 sub-categories defined by crossing natural vs. manmade with open vs. closed (Beach, Cavern, Street, and Livingroom). Within each face/scene sub-category, we selected 8 exemplars as dissimilar as possible to each other. Face dissimilarity was determined using the embeddings from the penultimate layer of the Dlib face model [87,88], which were further transformed to better reflect how humans perceive similarity between faces [89]. We applied k-means clustering to the embeddings of all faces for each face sub-category in the dataset. We then created eight clusters, and selected one representative exemplar from each cluster, whose embedding was closest to the gravitational center of the cluster, resulting in the selection of eight dissimilar representative exemplars within the sub-category. Scene dissimilarity was determined in a similar way but by using the embeddings from the output layer of GoogleNet pre-trained with Places365 datasets [90] (“imagePretrainedNetwork” function in MATLAB). After selecting all 32 stimuli, we scaled the lightness (Value in the HSV color space) separately for the face and scene stimulus sets so that the mean lightness was identical within each set. The face stimuli were blended and resized so that they fit within the boundaries of the scene stimuli (7.8 × 7.8 degrees of visual angle).

Since it was not feasible to present all 32 × 32 combinations of face and scene exemplars within a single scanning session, we selected 128 balanced subsets of combinations. Each combination was presented only once per session (128 trials in total in a session). Specifically, each face exemplar appeared four times per session, each time paired with a different scene exemplar from a different sub-category, and vice versa for scene exemplars. Half of the 128 combinations were presented in face-cued trials, and the other half in scene-cued trials. In the second session, the pairing between cues and stimuli was reversed, with all combinations presented again but in a new, randomized order. The third and fourth sessions repeated the same stimulus combinations and cues as the first and second sessions, respectively, but with the order of trials randomized again. Each scanning run showcased 16 out of 128 combinations, including all combinations of face and scene sub-categories (4 face sub-categories × 4 scene sub-categories = 16 trials).

To minimize eye movements in the scanner, we asked observers to focus their gaze on a fixation point [91], displayed at the center of the sample/test (i.e., face–scene blended stimuli).

### Task and procedure

Observers performed a delayed-match-to-sample task of either face or scene in the MRI scanner. At the beginning of each trial, a 100% predictive word cue (either “FACE” or “SCENE”) was presented in Arial font for 1s to indicate whether to remember a face or a scene in that trial (Figure 1A). Following that cue, a face-scene blended stimulus was presented as a “sample” stimulus, flashing on (0.8 s) and off (0.2 s) for a total of 9 cycles. The final (9th) blank was replaced with a noise mask to signal the end of the encoding phase, followed by 10s delay period (maintenance phase), during which only the fixation point was shown. The delay period was followed by the appearance of another face-scene blended image as a test stimulus, the duration of which was 2s. Observers judged whether the cued aspect (i.e., either the face or scene) of the test stimulus was the same or different from that of the sample stimulus. Thus, the dimension along which the attended aspect was manipulated and the dimension of the behavioral response were orthogonal by design, ensuring that the face-vs-scene coding axis was free from confounds related to decision- or response-related activity. To reduce regressors’ collinearity, the test image was omitted in a random half of the trials, in which no behavioral response was required. The contrast polarity reversal of the fixation point signalled the end of the trial, followed by an inter-trial interval (ITI), which was randomly chosen from 4, 5, 6, and 7s. Each functional run consists of 16 trials, with face-cued and scene-cued trials performed in equal numbers (8 trials each) and randomized order.

Out of eight trials requiring responses in each run, in two trials the face aspect changed, in two the scene aspect changed, in two both changed, and in two neither changed between the sample and test; within each type, one was face-cued and one scene-cued trial. In trials where they were intended to change, the test image was randomly selected from the *same* sub-category but from the entire dataset (rather than from the only eight exemplars, which served as the sample stimuli), to make the task moderately engaging.

### MRI data acquisition

Functional images were acquired using a T2*-weighted gradient-echo echo-planar imaging (EPI) sequence on a 7T Siemens MAGNETOM Terra scanner with a 32-channel head coil at the Center for Neuroscience Imaging Research. Acquisition parameters were as follows: repetition time (TR) = 1400 ms; echo time (TE) = 21.0 ms; flip angle = 61°; voxel size = 1.5 × 1.5 × 1.5 mm³; field of view (FOV) = 210 × 210 mm²; 92 axial slices; phase encoding direction = posterior to anterior; multiband acceleration factor = 4; GRAPPA acceleration factor = 2; phase partial Fourier = 6/8; bandwidth = 1984 Hz/pixel. Each functional run consisted of 319 volumes (including initial 5 volumes and final 10 volumes without any events), acquired in an interleaved slice acquisition order. Slice orientation was set obliquely to the AC-PC line (approximately -15° to axial). Each observer performed 8 functional runs in each of 4 scanning sessions, a total of 32 runs (due to scheduling constraints, Obs. 9 completed the final run of the third session in the fourth session.).

To correct for susceptibility-induced geometric distortions in the functional data, a pair of gradient-echo EPI volumes with reversed phase-encoding directions, one with anterior-to-posterior and one with posterior-to-anterior, was acquired before every functional run. These fieldmap images were acquired with the same pulse sequence as the functional scans, with each consisting of 4 volumes.

Structural T1-weighted images were acquired at the beginning of the first session (and additionally at the beginning of the third session for the last four observers). A 3D Magnetization-Prepared Two Rapid Acquisition Gradient Echoes (MP2RAGE) sequence was used with the following parameters: TR = 4500 ms; TE = 2.24 ms; inversion times (TI1/TI2) = 1000/3200 ms; flip angles = 4°/5°; voxel size = 0.7 × 0.7 × 0.7 mm³; FOV = 224 × 224 mm²; 320 sagittal slices; phase encoding direction = anterior to posterior; bandwidth = 200 Hz/pixel; acquisition time = 306 s. Automatic background denoising was applied, and the resulting image (UNI-DEN) was used as the T1w anatomical input for subsequent preprocessing in fMRIPrep.

### Preprocessing

MRI data were preprocessed using fMRIPrep 23.2.1 [92,93], which is built on Nipype 1.8.6 [94,95]. Many internal operations, particularly within the functional processing workflow, relied on Nilearn 0.10.2 [96].

### Fieldmaps

B0 nonuniformity maps (fieldmaps) were estimated from EPI references with opposite phase-encoding directions using FSL TOPUP v2203.2 [97] and applied to correct susceptibility distortions for each following functional run. For Obs. 2, due to a technical issue, the posterior-to-anterior reference for run 3 in session 1 was unavailable; in this case, the posterior-to-anterior reference from run 4 was used instead.

### Anatomical data

Each T1-weighted (T1w) image was corrected for intensity non-uniformity using N4BiasFieldCorrection (ANTs 2.5.0; [98, 99]) and skull-stripped using a Nipype implementation of the ANTs antsBrainExtraction.sh workflow with the OASIS30ANTs template. Brain tissue segmentation into cerebrospinal fluid (CSF), white matter (WM), and gray matter (GM) was performed on the brain-extracted T1w image using FSL FAST v2111.3 [100]. For observers with two T1w images, an anatomical T1w reference was generated by registering the INU-corrected images using mri_robust_template (FreeSurfer 7.3.2; [101]). Cortical surface reconstruction was performed with recon-all (FreeSurfer 7.3.2; [102]), and the brain mask was refined using a custom procedure based on Mindboggle to reconcile ANTs- and FreeSurfer-derived cortical segmentations [103]. Volume-based spatial normalization to MNI152NLin2009cAsym was performed using nonlinear registration with antsRegistration (ANTs 2.5.0), aligning brain-extracted T1w images to the ICBM 152 nonlinear asymmetric template version 2009c [104] accessed via TemplateFlow (v23.1.0) [105].

For one observer (Obs. 8), skull stripping failed within fMRIPrep; therefore, cortical surfaces were reconstructed using FreeSurfer after manual skull stripping of the T1w image prior to fMRIPrep processing.

### Functional data

For each BOLD run, the following preprocessing was performed using fMRIPrep. A reference volume was generated for motion correction, and head-motion parameters were estimated relative to this reference using FSL-MCFLIRT v2111.0 [106].

Susceptibility distortion correction was applied by aligning the estimated fieldmap to the EPI reference, which was subsequently coregistered to the T1w reference using boundary-based registration with six degrees of freedom (FreeSurfer bbregister; [107]). Framewise displacement was calculated for each functional run using both the formulations of Power et al. [108] and Jenkinson et al. [106], as implemented in Nipype.

Finally, BOLD time series were resampled onto the fsaverage surface (FreeSurfer format). All volumetric resamplings were performed in a single interpolation step by composing motion correction, susceptibility distortion correction, and coregistration. Volumetric resampling used cubic B-spline interpolation, and surface resampling was performed using FreeSurfer’s mri_vol2surf, followed by spherical resampling to fsaverage using FreeSurfer’s output *sphere.reg*.

### Quality control

We flagged two functional runs with poor preprocessed image quality. One deviated from the other runs of the same observer, possibly due to coregistration failure (Obs. 2, Session 1, Run 3), and the other exhibited severe striping artifacts in the inferior brain regions, likely caused by head motion during shimming (Obs. 5, Session 2, Run 4). Importantly, we confirmed that excluding these runs did not alter the subsequent group-level statistical results except where noted.

### General linear model

We fitted general linear models (GLMs) to the BOLD time courses per each vertex in the fsaverage space using the run_glm function of Nilearn (version 0.12.0). For each run, the encoding and maintenance phases were modeled with four separate regressors corresponding to face encoding (FE), scene encoding (SE), face maintenance (FM), and scene maintenance (SM) conditions, each of which was the 10-s box-car function convolved with the canonical SPM hemodynamic response function (HRF) (Figure S8). Note that the encoding phase included cue and mask in addition to the stimulus period (i.e., sample/blank) while the maintenance phase just included the 10s delay period. Responses to test images (appeared only on half of the trials) were modeled with another single regressor, irrespective of condition. Six motion parameters and polynomial drift terms up to the third order were included as nuisance regressors. When framewise displacement exceeded 0.5 mm, the signals for that TR were interpolated using neighboring TRs. This “runwise” GLM produced a beta matrix of size 4 (FE, SE, FM, and SM phases) × N (number of vertices) per run. For each run, the beta matrix was z-standardized within ROI by subtracting its mean and dividing by its standard deviation.

To increase the amount of training data for subsequent decoding analyses, we also estimated “trialwise” beta values using GLMsingle [109]. First, we upsampled the BOLD time series to a TR of 1 s [110], ensuring that event durations were integer multiples of the TR. Each event (FE, SE, FM, SM, and the test stimulus) was then modeled with a sequence of consecutive 1-s regressors spanning the entire event duration. GLMsingle selected the best-fit HRF for each vertex from a predefined library, identified nuisance regressors from pools of noise-dominant vertices, and finally applied ridge regression to stabilize single-trial beta estimates, with the degree of regularization determined by cross-validation [109]. Beta estimates were first obtained separately for each 1-s regressor and were then averaged across time points to yield a single beta for each event. This “trialwise” GLM produced a beta matrix of size 2 (encoding and maintenance phases) × N (number of vertices) per trial. Since one run has 16 trials, we obtained a beta matrix of size 32 × N per run, which was z-standardized within ROI prior to decoding analyses.

### Definition of ROIs

We examined the entire cerebral cortex, divided into 22 anatomically defined regions of interest (ROIs), which are combined subsets of the Human Connectome Project-MultiModal Parcellation (HCP-MMP) atlas exactly following the grouping strategy in Glasser et al. [34] (Figure S2). In addition, we also used a more fine-grained definition of ROIs, namely the original 180-region HCP-MMP atlas, for visualization and as exploratory analyses. We analyzed corresponding regions in the left and right hemispheres as a single ROI.

### Rotation angle of the coding axis

To quantify the geometric rotation from the encoding to the maintenance phases of the working memory, we measured the angle between their corresponding face–scene coding axes (defined by runwise GLM beta patterns for face-cued vs. scene-cued trials). There were a total of 32 runs, each resulting in a 1×N (vertices) face-scene coding axis for the encoding and another for the maintenance phase within each ROI. We averaged the coding axes across runs and computed the angle between the coding axes of the encoding and maintenance phases:

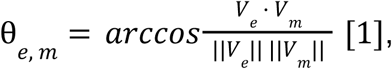

where 𝑉_𝑒_ and 𝑉_𝑚_ denote the run-averaged face-vs-scene coding axes for encoding and maintenance phase, respectively. To obtain the distribution of θ_𝑒, 𝑚_, we used bootstrap resampling. For each ROI, starting from the 32 (number of runs) × *N* matrix for runwise coding axes, we generated 10,000 bootstrap replicates by resampling new 32 rows with replacement and averaging across the resampled rows, thereby yielding one bootstrapped estimate of the coding axis each for encoding and maintenance phase per replicate. We then calculated the angle between the two axes for every replicate.

The angle between two parallel axes can even appear as non-zero due to noise, with larger noise leading to larger deviations from zero. Thus, to evaluate whether two axes are truly rotated (i.e., non-parallel), we compared their observed angle (i.e., the distribution obtained above) to the angle expected to arise purely from noise, even if the axes are parallel (i.e., null hypothesis of parallelism). This expected rotation caused purely by noise (hereafter referred to as noise floor) follows Equation 2 (see Supplementary Information for the derivation in detail):

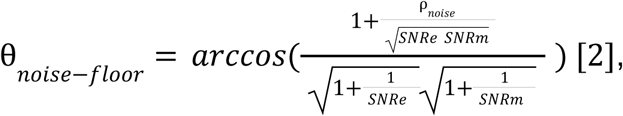

𝑆𝑁𝑅𝑒 and 𝑆𝑁𝑅𝑚 are the signal-to-noise ratios in the coding axis of the encoding and maintenance phase, respectively. Lower SNR (i.e., greater noise levels) raises the noise floor of the angle, as indicated by the equation above. This means that even if the axes are parallel, for larger noise levels, the observed angle could deviate further from 0.

ρ_𝑛𝑜𝑖𝑠𝑒_ indicates the cross-noise correlation between encoding and maintenance phase coding axes. Given that encoding and maintenance phases both occurred on the same trials (see Figure 1A), we considered the possibility that their noise components might be correlated. Correlated noise could make the axes appear more parallel, thus it would lower the noise floor, as indicated by the equation above.

The SNR terms are estimated as follows:

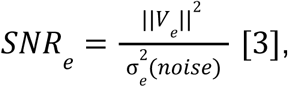

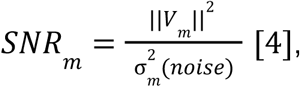

where ||𝑉_𝑒_||^2^ and ||𝑉_𝑚_|^2^| are the true squared L2 norms of the axis for encoding and maintenance phase, respectively, approximated as follows from the observations.

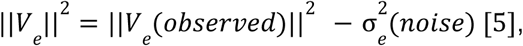

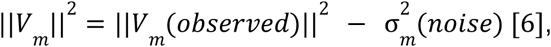

𝑉_𝑒_ (𝑜𝑏𝑠𝑒𝑟𝑣𝑒𝑑) and 𝑉𝑚 (𝑜𝑏𝑠𝑒𝑟𝑣𝑒𝑑) indicate the observed coding axes for encoding and maintenance phase (averaged across runs), respectively.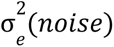 and 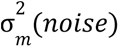 denote the noise variances. Noise variances were estimated via the bootstrap resampling described above. For each dimension (vertex), we computed the variance across bootstrap replicates and then summed these variances across dimensions to obtain the total noise variance.

Note that when noise is overwhelming, the above estimations of ||𝑉||^2^ could become negative, thus resulting in a negative SNR. To avoid introducing negative SNRs in Equation 3, which would make the cosine undefined, we clipped||𝑉||^2^ to 0. This results in a noise floor of 90 degrees, suggesting that in such a noise level, even parallel axes are expected to appear orthogonal on average (in this clipped case, if no true rotation exists, the actual angle would still be distributed around 90 degrees, so the z-value would be around zero).

The cross-noise correlation is defined as follows:

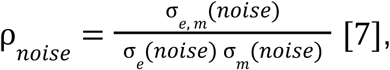

where σ_𝑒, 𝑚_ (𝑛𝑜𝑖𝑠𝑒) is the cross-noise covariance estimated as the covariance across bootstrap replicates within each dimension and then summed over dimensions. It is noteworthy that cross-noise correlation was generally small, so this term had little impact on the noise floor.

Finally, z-score of *rotation* was obtained per ROI and per observer by comparing the distribution of θ𝑒, 𝑚 with θ𝑛𝑜𝑖𝑠𝑒−𝑓𝑙𝑜𝑜𝑟 which, as discussed above, is the estimated mean of the null hypothesis (i.e., observed angle even if the axes are parallel):

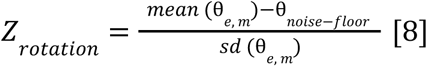

### Multivariate pattern decoding

#### Decoding face vs. scene

To assess whether visual content information was represented in each ROI, we trained linear decoders to classify contents (face vs. scene) irrespective of phases. We also repeated the same analysis separately for the encoding and maintenance phases.

#### Exclusive-or (XOR) decoding

More importantly, we also trained decoders to classify exclusive combinations (encoded face and maintained scene vs. encoded scene and maintained face). Successful classification in this so-called exclusive-or (XOR) decoding in an ROI would indicate that the two factorized variables (phase and content) are entangled within a stable three-dimensional representational subspace in that ROI [15].

#### Cross-Condition Generalization Performance (CCGP)

To assess the generalizability of a hyperplane distinguishing face–scene dichotomy across phases, we also measured cross-condition generalization performance (CCGP; see [15,48]). Specifically, we defined CCGP as the average of cross-decoding accuracy in both directions (trained a classifier to distinguish face vs. scene only in the encoding phase and tested it in the maintenance phase and vice versa).

#### Decoding analysis procedures

All the abovementioned decoding accuracies were assessed using 8-fold cross-validation on trial-wise GLM beta patterns, z-standardized within each run. To prevent information leakage, all data from the same run was assigned to the same fold (using GroupKFold in Scikit-learn), which also helped balance conditions across folds. For example, to examine CCGP, decoders were trained to classify face vs scene using data from one phase (e.g., encoding) in seven folds and tested on data from the other phase (e.g., maintenance) in the held-out fold. For each iteration of cross-validation, PCA was applied to the training data (retaining components that explain 95% of the variance), followed by training a logistic regression model (using Scikit-learn library). Test data was projected onto the same PC space, and the model prediction accuracy was calculated and averaged across folds. To assess the statistical significance of the decoding accuracy, a null distribution was generated by permuting condition labels within each run and repeating the entire cross-validation procedure 1,000 times [111].

#### Tradeoff between geometric stability and flexibility

Theoretically, in factorized low-dimensional geometries, CCGP approaches perfect, while XOR performance remains at chance (Figure 1B). Conversely, in entangled high-dimensional geometries, XOR performance exceeds chance, but CCGP decreases (Figure 1D). This pattern reflects a tradeoff between flexibility (indicated by above-chance XOR decodability) and stability (indicated by above-chance CCGP) in general information-coding principles. In an exploratory analysis, we quantified this tradeoff per ROI as:

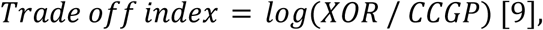

where the logarithmic transformation ensures that the metric is symmetric around a balanced tradeoff (index = 0). We used this index as a convenient summary of the geometric flexibility–stability tradeoff, while noting that an alternative, potentially fairer tradeoff measure was explored in simulations and yielded similar results overall (see Supplementary Information and Figures S9, S10, S11, and S12).

### Statistical analyses

We performed group-level statistical tests based on our eight observers. For the angle rotation, group-level significance was evaluated using one-tailed one-sample t-tests (against zero) on the individual z-values (see *Rotation angle of the coding axis* section). For the decoding analysis, we similarly reported the results of one-tailed one-sample t-tests (against zero) performed on the individual z-values of decoding performances. We also note that the significance patterns were unchanged when logit-transformed decoding accuracies were tested instead. All p-values were corrected for multiple comparisons across all ROIs using the false discovery rate (FDR) [112].

We also assessed whether significance at the individual level was universal [35]. Based on the pre-registered sensitivity of tests (1 – β = .9), we considered the significance as universal if at least 5 (out of 8) observers showed significance at the individual level, whereas we considered it as sporadic if at least 6 (out of 8) observers showed non-significance.

## Data availability

Anonymized MRI data are publicly available on OpenNeuro (https://openneuro.org/crn/reviewer/eyJhbGciOiJIUzI1NiIsInR5cCI6IkpXVCJ9.eyJzd WIiOiJmZTA5NjVlMi1kMTcxLTRkOWUtYmMzOS0wNTY5M2I0MjY1YjAiLCJlbWFpb CI6InJldmlld2VyQG9wZW5uZXVyby5vcmciLCJwcm92aWRlciI6Ik9wZW5OZXVybyIs Im5hbWUiOiJBbm9ueW1vdXMgUmV2aWV3ZXIiLCJhZG1pbiI6ZmFsc2UsInNjb3Blc yI6WyJkYXRhc2V0OnJldmlld2VyIl0sImRhdGFzZXQiOiJkczAwNzA4MiIsImlhdCI6MT c3OTI2MzM2MSwiZXhwIjoxODEwNzk5MzYxfQ.-BnPPNm4uGLOa24RSWTIWAUM Ng9YDcAyGJjFeaKUU28).

GLM outputs (beta values for each vertex) are publicly available on Zenodo (for runwise beta, https://zenodo.org/records/18203411?preview=1&token=eyJhbGciOiJIUzUxMiJ9.eyJ pZCI6IjE0Nzc4ZGE2LTI5ZTktNDU3Mi1hY2Q1LWI4YzA5ODZmMTFhZiIsImRhdGEiOnt9LCJyYW5kb20iOiJhNmM3NTM1NzMyMGQwYzE5ZWZlNDJhMDI4NjUyY2Y1Zi

J9.YQ85cji6C5VyhbFqkhzllUoPXDcyNJ7-isLknMBwG7XxoqL2v_7ZvlW3JcewsbnZU J6pfOuf407tbBlP-MbLDg; for single-trial beta, https://zenodo.org/records/18203430?preview=1&token=eyJhbGciOiJIUzUxMiJ9.eyJ pZCI6ImVjMjA3YTJkLTgzYWMtNGVkMy1hZmUxLTgwYThjNzVmNzAwZSIsImRhdG EiOnt9LCJyYW5kb20iOiI4M2Q2NGRhM2YxMDY1YThiOGMyZjBiYTgwNDA4MDgx YyJ9.lZ04iekodhCsCqtMuHgyTGCFDF9gfN4MAhmJGq5xAgtR0-hDCV9TAoEnD6F Do7aSaFGbojgP61YBV-75Yi_1Kg and https://zenodo.org/records/18204325?preview=1&token=eyJhbGciOiJIUzUxMiJ9.eyJ pZCI6ImM0ZDdmOGZkLTMwMTAtNDY5Yy05OTZlLTJhYTU3Y2UyMmExNSIsImRh dGEiOnt9LCJyYW5kb20iOiIzNDA0Mzc2MDYxYWU0MzU3ZTJmYzE4OWFhNWQzNDhlZiJ9.EaAghc7_mNN9xu4KuWqK3ZIwhQ4nd856L1YXO5jTBK7FBLqK8WSSgw 4IX2L7hZSXmgv8Udk421dZGMY0I-s7hA).

Behavioral data are publicly available on GitHub (https://github.com/Tomnakam/geom-wm-dynamics).

## Code availability

All original code used for the experiments and analyses is publicly available on GitHub (https://github.com/Tomnakam/geom-wm-dynamics).

## Acknowledgements

T.N. is supported by JSPS KAKENHI Grant Numbers 24KJ0233, 25K18957, and 25K00896 in Japan. H.L. is supported by the Institute for Basic Science

(IBS-R015-D2) in the Republic of Korea. We thank Dr. Joshua Bradley Chan Tan for helpful comments on the manuscript.

## Author contributions

Conceptualization, T.N., H.L., and A.M.; methodology, T.N., M.Y., K.K., H.L., and A.M.; software, T.N.; validation, T.N., K.K., and A.M.; formal analysis: T.N., and A.M.; investigation: T.N.; resources: H.L.; data curation: T.N., K.K.; writing - original draft, T.N.; writing - review & editing, T.N., M.Y., K.K., H.L., and A.M.; visualization, T.N., and A.M.; supervision: H.L., M.Y., and A.M.; project administration: T.N.; funding acquisition: T.N., and H.L.

## Competing interests

The authors declared no competing interests.

## Notes

### Competing Interest Statement

The authors have declared no competing interest.

### Summary of Updates

In this revision, we changed the conceptual framing of the manuscript from a perception-versus-working-memory comparison to an investigation of representational differences between the encoding and maintenance phases of working memory.

