## Supplementary Information for "Representational Geometries Across Visual Working Memory Encoding and Maintenance"

### Derivation of Equation 2

The full expression for the cosine similarity between the coding axis for the encoding phase $V_e$ and the maintenance phase $V_m$, incorporating noise terms, $\epsilon_e$ and $\epsilon_m$, is:

$$cos(V_e + \epsilon_e, V_m + \epsilon_m) = \frac{V_e \bullet V_m + \epsilon_e \bullet V_m + V_e \bullet \epsilon_m + \epsilon_e \bullet \epsilon_m}{\sqrt{||V_e||^2 + 2V_e \bullet \epsilon_e + \epsilon_e \bullet \epsilon_e}\sqrt{||V_m||^2 + 2V_m \bullet \epsilon_m + \epsilon_m \bullet \epsilon_m}}$$

Under the assumption that signal and noise are independent, and the noises have a zero mean, the expression reduces to:

$$cos(V_e + \epsilon_e, V_m + \epsilon_m) = \frac{V_e \bullet V_m + \sigma_{e,m}(noise)}{\sqrt{||V_e||^2 + \sigma_e^2(noise)}\sqrt{||V_m||^2 + \sigma_m^2(noise)}}$$

$$= \frac{V_e \bullet V_m + \sigma_{e,m}(noise)}{||V_e||\,||V_m||\sqrt{1 + \frac{\sigma_e^2(noise)}{||V_e||^2}}\sqrt{1 + \frac{\sigma_m^2(noise)}{||V_m||^2}}}$$

$$= \frac{\frac{V_e \bullet V_m}{||V_e||\,||V_m||} + \frac{\sigma_{e,m}(noise)}{||V_e||\,||V_m||}}{\sqrt{1 + \frac{\sigma_e^2(noise)}{||V_e||^2}}\sqrt{1 + \frac{\sigma_m^2(noise)}{||V_m||^2}}}$$

Here, $\sigma_e^2(noise)$, and $\sigma_m^2(noise)$ represent the noise variances of coding axis for the encoding phase and maintenance phase, respectively, and $\sigma_{e,m}(noise)$ indicate their noise covariance. Under the null hypothesis of parallelism, the cosine value between $V_e$ and $V_m$ is 1.

$$cos(V_e + \epsilon_e, V_m + \epsilon_m) = \frac{1 + \frac{\sigma_{e,m}(noise)}{||V_e||\,||V_m||}}{\sqrt{1 + \frac{\sigma_e^2(noise)}{||V_e||^2}}\sqrt{1 + \frac{\sigma_m^2(noise)}{||V_m||^2}}}$$

Then, for a better intuitive understanding of the equation, we rewrite it using the SNR notion.

$$cos(V_e + \epsilon_e, V_m + \epsilon_m) = \frac{1 + \frac{\frac{\sigma_{e,m}(noise)}{\sigma_e(noise)\,\sigma_m(noise)}}{\frac{||V_e||}{\sigma_e(noise)}\,\frac{||V_m||}{\sigma_m(noise)}}}{\sqrt{1+\frac{1}{\frac{||V_e||^2}{\sigma_e^2(noise)}}}\sqrt{1+\frac{1}{\frac{||V_m||^2}{\sigma_m^2(noise)}}}}$$

$$= \frac{1+\frac{\rho_{noise}}{\sqrt{SNRe\ SNRm}}}{\sqrt{1+\frac{1}{SNRe}}\sqrt{1+\frac{1}{SNRm}}}$$

#### Supplementary figures

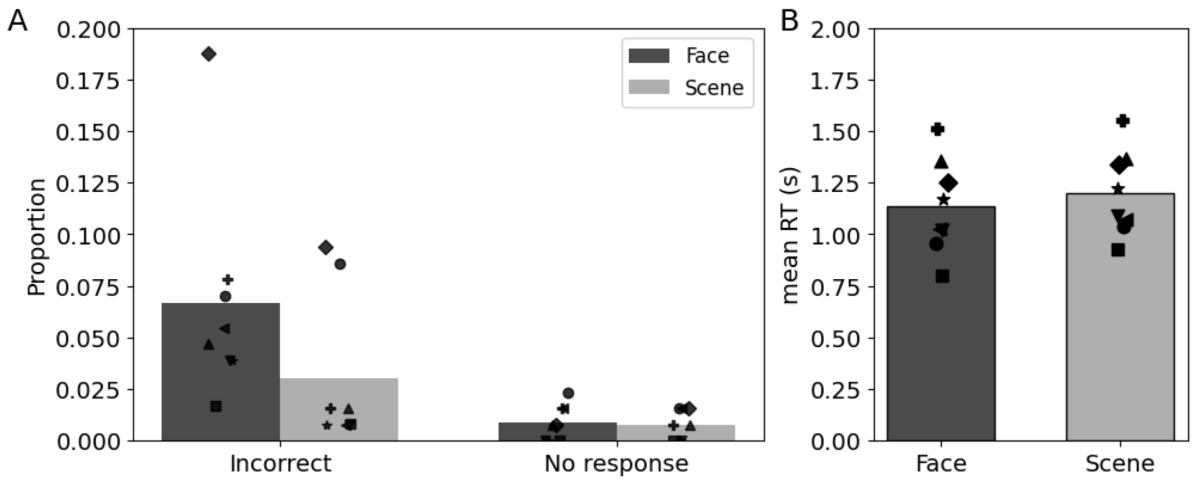


***Figure S1. Behavioral data.*** *(A) Proportion of trials (among those including the Test phase) that resulted in incorrect responses and no responses. The mean correct response rate was 94.3% and the proportion of no-response trials was low (mean = 0.8%). At the group level, correct response rates for the scene-cued trials were significantly higher than for the face-cued trials (paired t-test: t(7) = 3.44, p = .011). (B) Geometric mean reaction time (RT) for correct trials in each observer. In contrast to correct response rates, RTs were significantly shorter for the face-cued trials (paired t-test on mean log-RTs: t(7) = 4.03, p = .005). Bar lengths indicate the inter-observer mean. Note that we excluded two runs (1st run in the 1st and 3rd sessions) for Obs. 2 only from behavioral analyses because the responses were not properly recorded due to technical issues.*

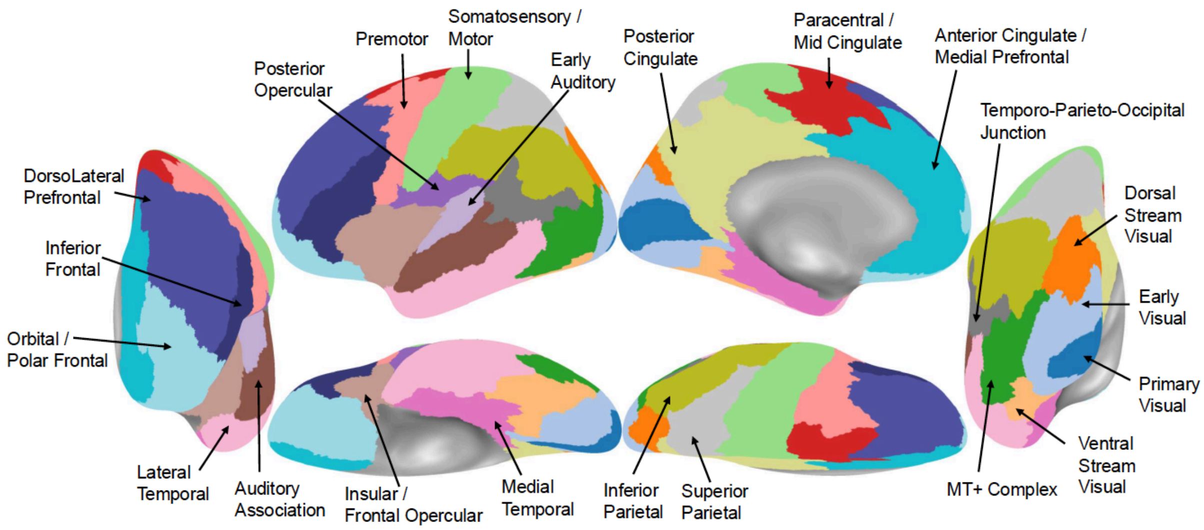


***Figure S2. Definition of the regions of interest (ROIs) used for the main analyses.*** *Different colored regions indicate each of the 22 coarse-grained ROIs defined by Glasser et al. [s1].*

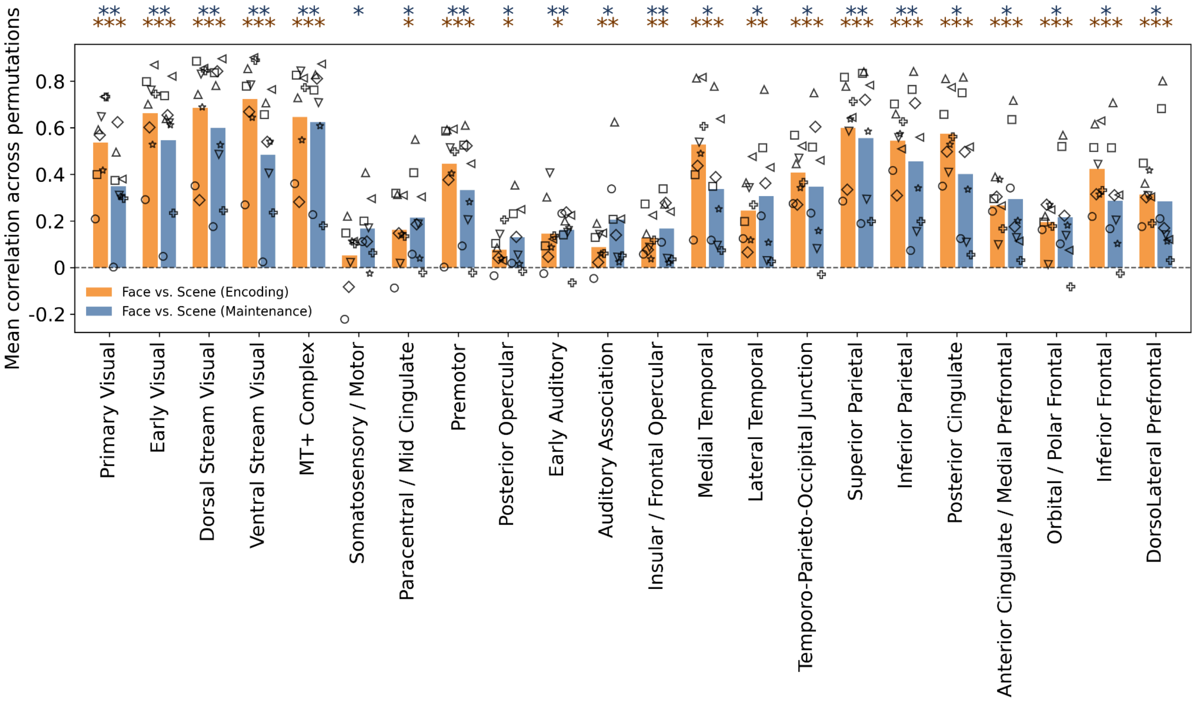


***Figure S3. Test-retest reliability of face-vs-scene tuning.*** *Each marker indicates each observer, showing the mean correlation across 10,000 split-half permutations. For each permutation, runs were randomly split into two halves, and the Pearson correlation was computed between the multivertex activation patterns (face-scene contrast) of the two halves across all vertices within each ROI. Asterisks indicate group-level statistical significance, assessed by one-sample one-tailed t-test (*** q < .001, ** q < .01, * q < .05; FDR-corrected among all ROIs). Face-vs-scene activity pattern was reliable across different runs, particularly in visual and most association areas (temporal, parietal, prefrontal cortex) both for encoding and maintenance phases. Such reliable activity patterns provide a basis for the subsequent analysis regarding the angle between face-vs-scene coding axes.*

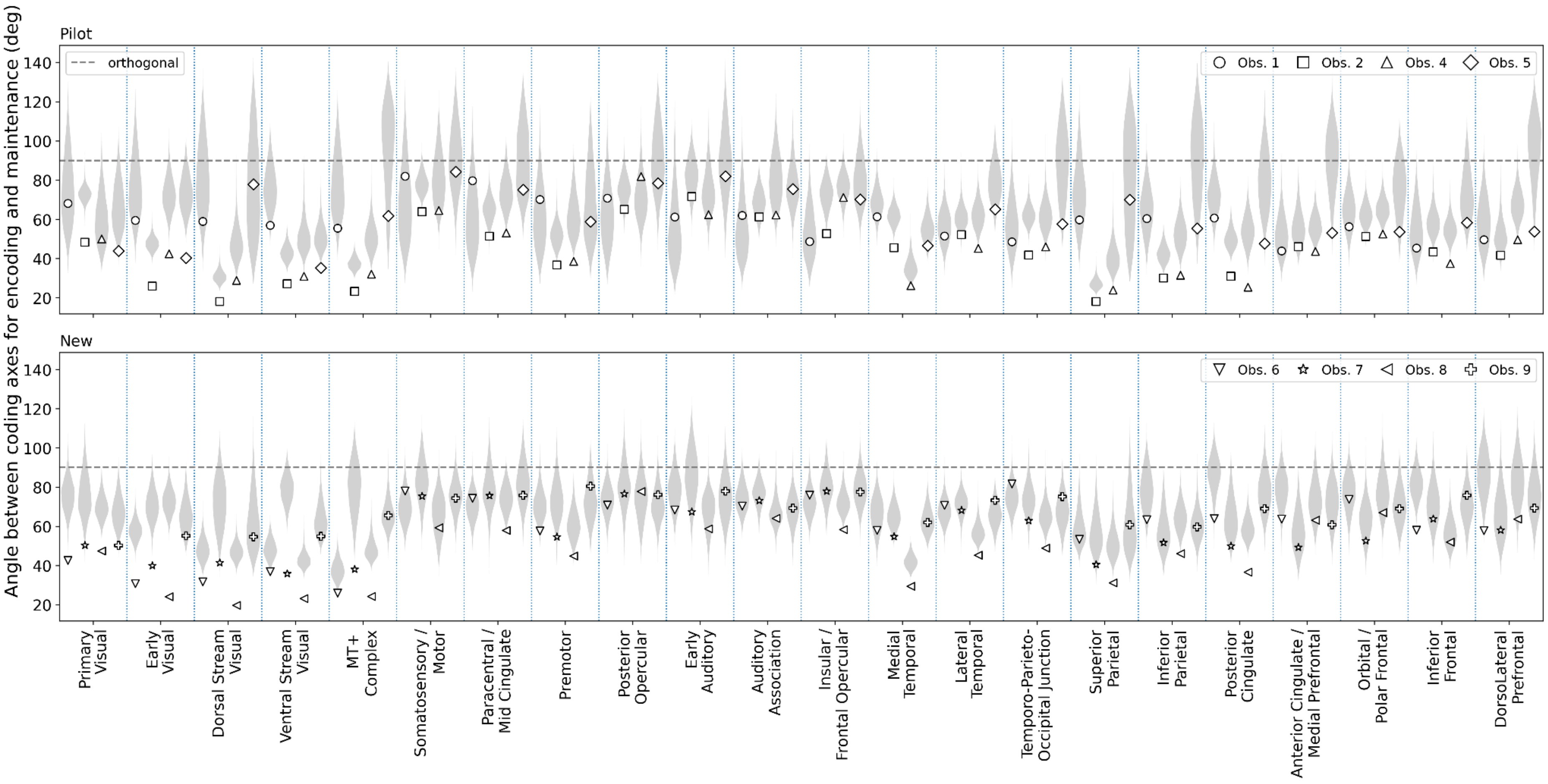


***Figure S4. Individual results of angle analysis.*** *Bootstrapped distribution of angle between face-vs-scene coding axes during encoding and maintenance phases was shown along with its noise floor. Different marker shapes indicate the noise floor from different observers.*

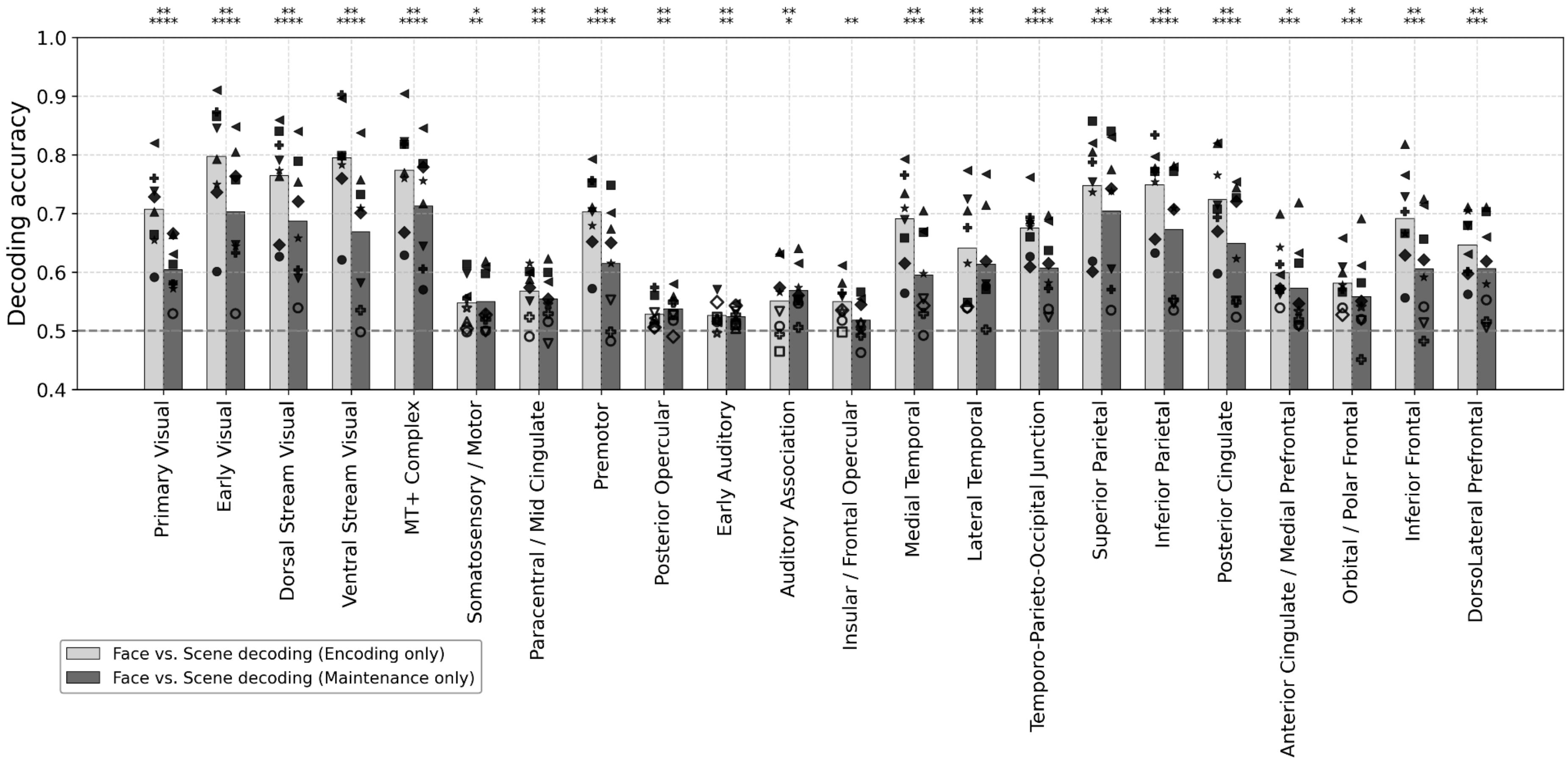


***Figure S5. Decoding accuracy for face vs. scene assessed separately within the encoding and maintenance phases.*** *Asterisks denote the significance of the group-level one-sample t-test (one-tailed) on z-transformed accuracy, FDR-corrected across all ROIs (**** q < .0001, *** q < .001, ** q < .01, * q < .05). These results are consistent with the test-retest reliability results for face-scene activity pattern shown in Figure S3.*

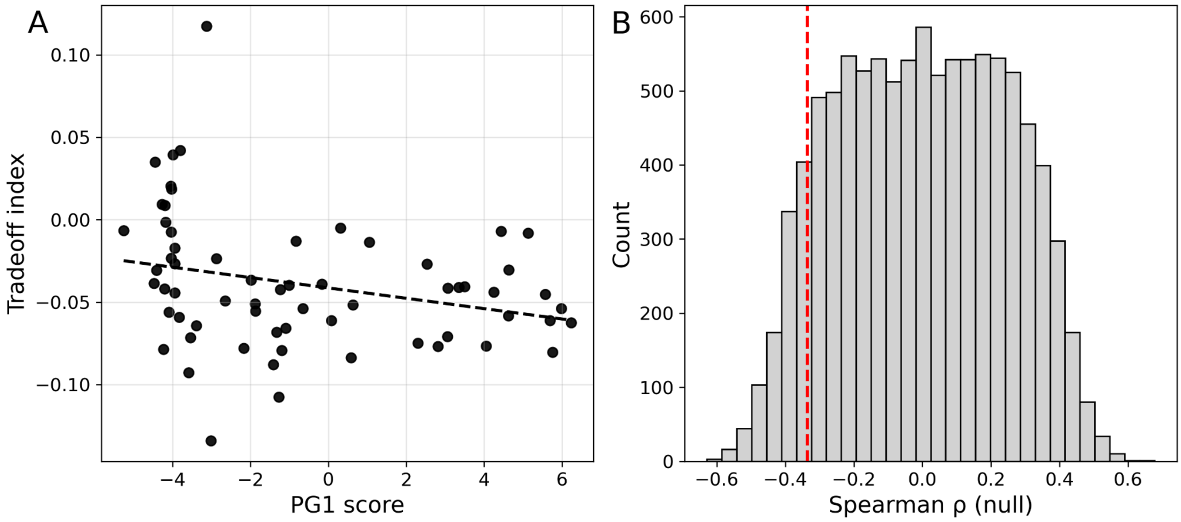


***Figure S6. Relationship between the tradeoff index and the functional gradient.*** *(A) Tradeoff index plotted against the 1st principal gradient scores from Margulies et al. [s2] for the ROIs selected using the same criteria as in Figure 4B. The Spearman correlation was significantly negative when tested against a null hypothesis assuming fully random spatial structure ($\rho = -.336$, two-sided $p = .007$). (B) Null distribution of Spearman $\rho$ obtained while preserving spatial autocorrelation [s3, s4]. Under this more stringent null model, the observed correlation was not significant (two-sided $p = .188$). The red vertical line indicates the observed correlation coefficient.*

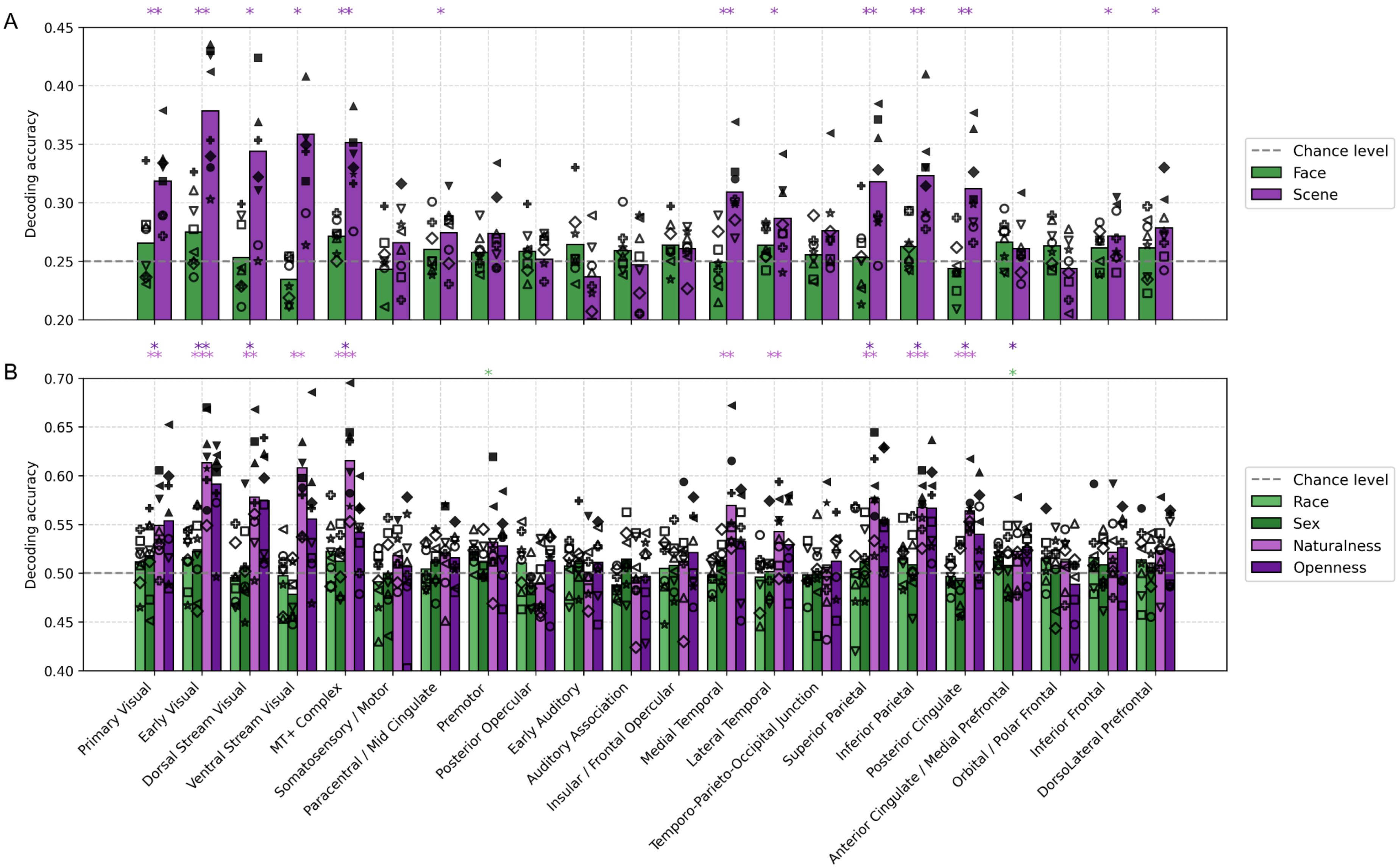
A
B
Decoding accuracy
Chance level
Face
Scene
Race
Sex
Naturalness
Openness
Primary Visual
Early Visual
Dorsal Stream Visual
Ventral Stream Visual
MT+ Complex
Somatosensory / Motor
Paracentral / Mid Cingulate
Premotor
Posterior Opercular
Early Auditory
Auditory Association
Insular / Frontal Opercular
Medial Temporal
Lateral Temporal
Temporo-Parieto-Occipital Junction
Superior Parietal
Inferior Parietal
Posterior Cingulate
Anterior Cingulate / Medial Prefrontal
Orbital / Polar Frontal
Inferior Frontal
DorsoLateral Prefrontal

***Figure S7. Decoding results for specific sub-categories.*** *(A) Decoding accuracy among the four face sub-categories within face-cued trials (green) and among the four scene sub-categories within scene-cued trials (purple). (B) Decoding accuracy for black vs. white (light green) and female vs. male (deep green) within face-cued trials, and for natural vs. manmade (light purple) and open vs. closed (deep purple) within scene-cued trials. All decoders were trained and tested irrespective of working memory phases. Asterisks denote the significance of the group-level one-sample t-test (one-tailed) on z-transformed accuracy, FDR-corrected across all ROIs (*** q < .001, ** q < .01, * q < .05). Overall, beyond face-vs-scene decoding, we successfully decoded scene sub-categories from the entire visual cortex as well as from the entire parietal cortex (superior parietal, inferior parietal, and posterior cingulate cortex), medial/lateral temporal cortex and only lateral parts of prefrontal cortex (inferior frontal and dorsolateral prefrontal cortex). In contrast, we could not reliably decode face sub-categories from any ROI. Furthermore, from similar areas, we were able to decode scene sub-category dichotomies (Figure 7B). The most decodable dichotomy was naturalness (natural vs. manmade scenes), which was decoded from the entire visual cortices, medial/lateral temporal cortex, and from the entire parietal cortices (superior parietal and inferior parietal). The second most decodable dichotomy was openness (open vs. closed scenes), which was decoded from some of the visual, parietal, and prefrontal cortices. Face subcategory dichotomy (only the race) was decoded only from the premotor and anterior cingulate/medial prefrontal cortex. The difference in sub-category decodability between faces and scenes may reflect the greater visual diversity of scenes compared to faces. In any case, these results confirm that, at least in the visual and some association (temporal, parietal, and lateral prefrontal) cortices, each scene sub-category is differentially represented and thus the rotation of the face-vs-scene coding axes of interest cannot be solely explained by language processing regarding the cue or by differences in the spatial extent of the attentional window [s5]. When assessed using logit-transformed accuracy, face decodability in MT+ complex became significant, whereas openness decodability in the primary visual cortex and posterior cingulate cortex became non-significant. Moreover, when the runs with poor image quality were excluded, scene decodability in the insular/frontal opercular cortex and naturalness decodability in the premotor cortex became significant, whereas openness decodability in the primary visual cortex and race decodability in the premotor cortex became non-significant.*

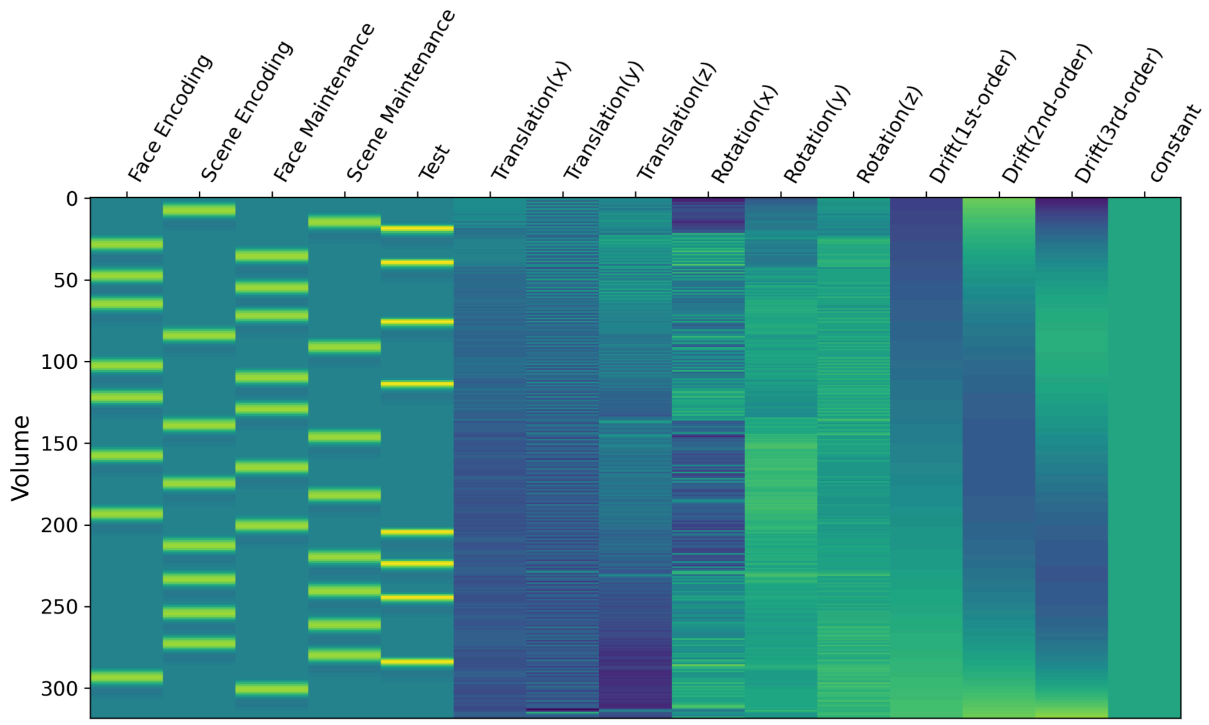


***Figure S8. Design matrix for the runwise GLM.*** *Each column indicates a regressor, and each row indicates a volume. The leftmost four regressors correspond to each condition of interest. The 5th regressor from the left corresponds to the test stimuli. 6th to 11th regressors are motion parameters derived during preprocessing. 12th to 14th regressors model slow polynomial drift (up to 3rd order). The rightmost regressor models the intercept for this run. One example run is shown here.*

#### Simulation of the tradeoff

CCGP is high for factorized, low-dimensional representational geometries, whereas XOR accuracy is high for high-dimensional, entangled geometries. Thus, CCGP and XOR should theoretically be in a tradeoff relationship. However, when we simply plot empirical CCGP accuracy against XOR accuracy across ROIs, they instead show a positive correlation (Figure S9A). This counterintuitive pattern can be explained by differences in functional SNR across regions (high SNR raises decoding accuracy overall). Here, we illustrate how SNR influences the apparent relationship between CCGP and XOR using simulations based on a simplified toy model and test the robustness of the results regarding tradeoff even after SNR effects are better accounted for.

In our simulation, we assumed three dimensions because four conditions can be fully represented in a three-dimensional space, and for simplicity we assumed isotropic Gaussian noise. We began with perfectly factorized ("rectangular") geometry that maximizes CCGP while rendering XOR decoding chance-level (Figure 1B). In this configuration, mean of the activation patterns for each of the four conditions lie on the same two-dimensional plane, and all observed responses can be generated by a model characterized by three parameters ($S_{E-M}$: signal strength along the encoding-maintenance axis, $S_{F-S}$: signal strength along the face-scene axis, $\sigma_{noise}$: unit variance Gaussian noise). Starting from this perfectly factorized geometry, we rotated the memory axis stepwise (by 5 degrees) in a direction orthogonal to the initial two-dimensional plane until the encoding and maintenance axis became orthogonal to each other. This orthogonal geometry represents the opposite end of the tradeoff continuum, supporting maximal XOR decodability while disabling CCGP (Figure 1D).

We found that under the assumption of independent noise, $S_{E-M}$ does not change the simulation results as long as we had enough trials. Therefore, here we report results obtained by fixing the $S_{E-M}$ as 2 (twice the $\sigma_{noise}$). When we plot XOR against CCGP, as the maintenance axis is rotated more, the point moves along the concave. Importantly, $S_{F-S}$ moves the curve toward upright (different colored curves in Figure S10A), which illustrates the fact that even if angle is the same, higher SNR produces higher XOR and higher CCGP simultaneously, which could explain the apparent positive relationship between them across different ROIs (Figure S9A).

To eliminate the effect of SNR, we normalized the measures as follows:

$$normalized\ CCGP = \frac{\Phi^{-1}(ACC_{CCGP})}{\Phi^{-1}(ACC_{FvsS})},\ normalized\ XOR = \frac{\Phi^{-1}(ACC_{XOR})}{\Phi^{-1}(ACC_{FvsS})}\ [S1],$$

where $ACC_{FvsS}$, $ACC_{XOR}$, $ACC_{CCGP}$ denote the decoding accuracies for the face-vs-scene dichotomy, XOR dichotomy and CCGP, respectively, and $\Phi^{-1}$ indicates the probit transformation. Probit normalization is justified under the assumption of Gaussian noise. Our simulations confirmed that the relationship between normalized CCGP and normalized XOR remains stable across different SNR levels (Figure S10B). In the empirical data, normalized CCGP and normalized XOR also showed a negative relationship (Figure S9B), although their values do not exactly follow the simulated trajectory (Figure S10B), because, in the real brain, unlike in the simulation, noise structure is likely more complex, and the vertex space is also higher-dimensional.

Given the improved normalization strategies, we can quantify the tradeoff for each region as the difference between the probit ratios, as follows:

$$Tradeoff\ index = normalized\ XOR - normalized\ CCGP\ [s2],$$

We confirmed that the tradeoff index increased monotonically with rotation angle and, importantly, that this relationship was largely independent of SNR (Figure S10D). It is noteworthy that the simpler definition of the tradeoff index, used in Figure 4, likewise exhibited a monotonic increase with angle (Figure S10C). Although SNR independence was not perfect, deviations were primarily observed in cases of poor SNR, which were largely excluded from the main analyses (see Figure 4), as well as at extreme rotation angles (0° or 90°), which were unlikely for most ROIs in the empirical data (see Figure S4).

Even when the effects of SNR were more precisely accounted for using the index, described in Equation S2, we further confirmed that the overall empirical results remained consistent: early sensory regions favored flexibility whereas association cortex favored stability (Figure S11; cf. Figure 4). The correlation between first functional gradient (unimodal-to-transmodal) [s2] and tradeoff index was slightly stronger, but still it was marginally significant ($p = .056$) when considering the spatial autocorrelation in the null distribution (Figure S12). However, given that this alignment with functional gradient was not statistically robust across indices (cf., Figure S6), this finding should be interpreted with caution and remains inconclusive.

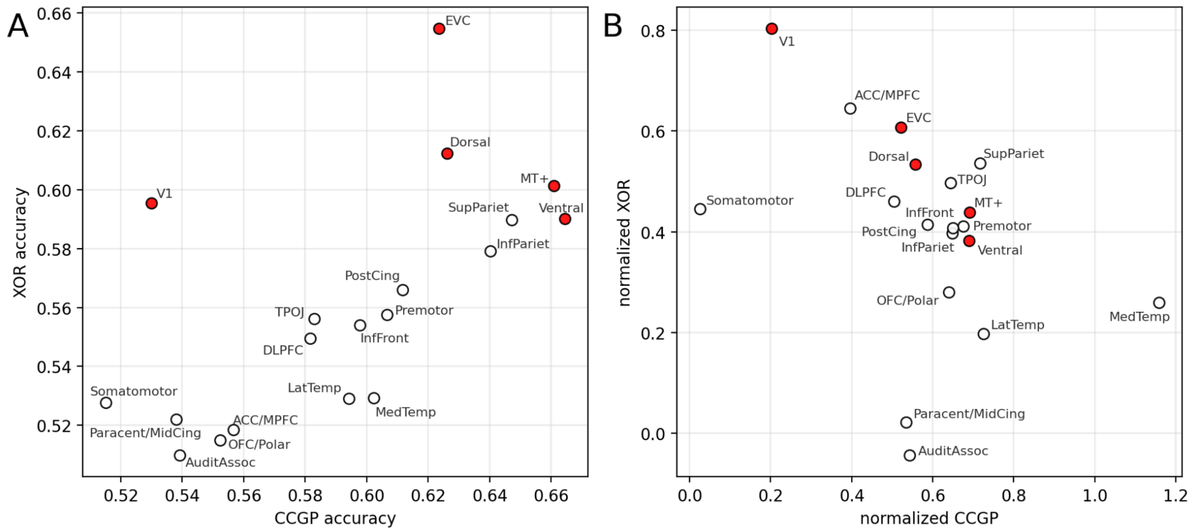


***Figure S9. The empirical relationship between CCGP and XOR.*** *(A) The relationship between interobserver mean of CCGP accuracy and XOR accuracy. (B) The relationship between interobserver mean of normalized CCGP and normalized XOR. Visual areas are highlighted in red.*

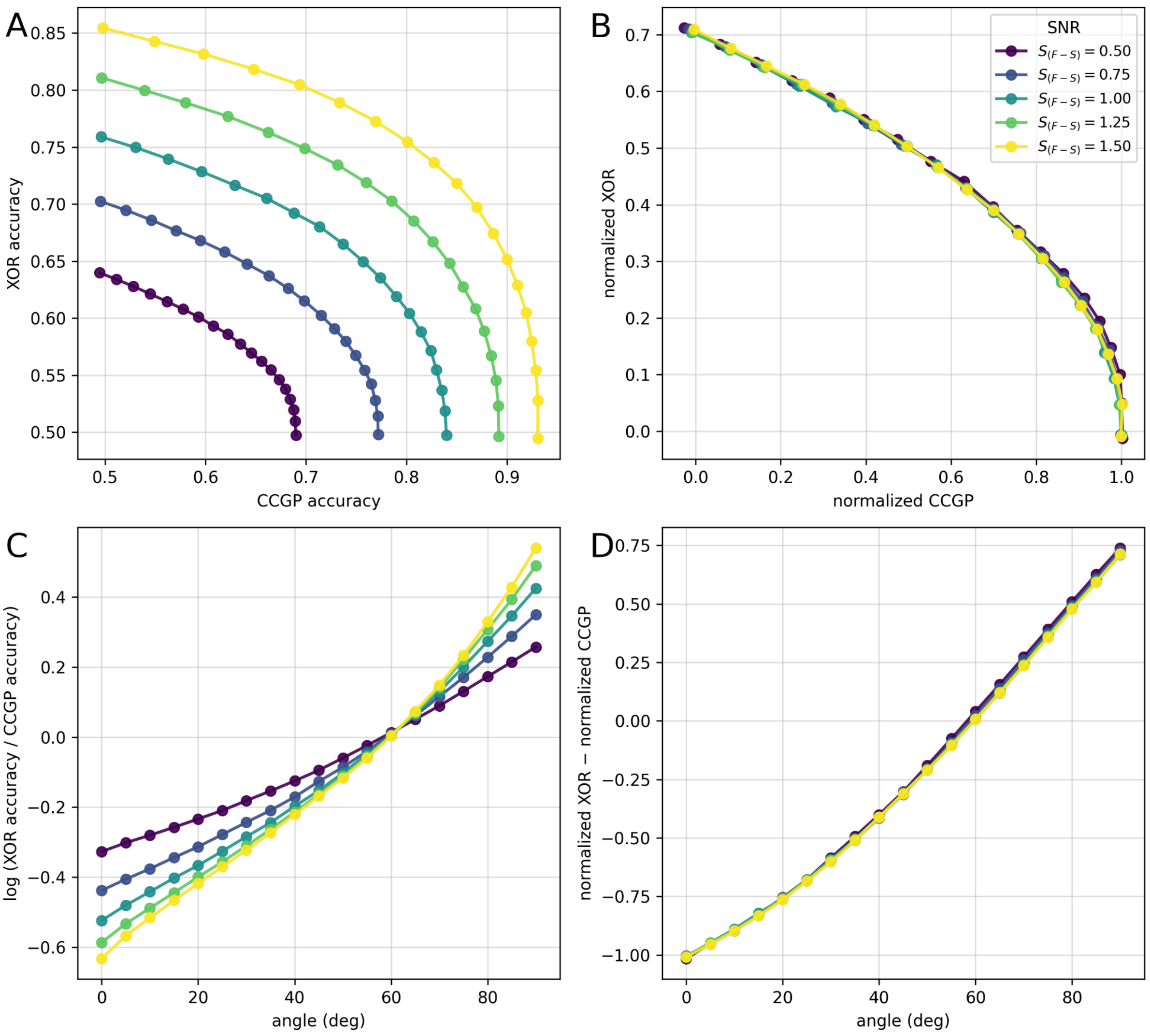


***Figure S10. Simulation for SNR-independent index of tradeoff.*** *(A) The simulated relationship between CCGP accuracy and XOR accuracy. (B) The simulated relationship between normalized CCGP and normalized XOR. Different colors indicate different levels of SNR. As the encoding and maintenance axes became increasingly orthogonal, points traversed the curve from the bottom right (maximal CCGP with chance-level XOR) toward the top left (chance-level CCGP with maximal XOR), with points sampled at 5° rotation increments. (C) The simulated relationship between rotation angle and tradeoff index defined in Equation 9. (D) The simulated relationship between rotation angle and tradeoff index defined in Equation S2.*

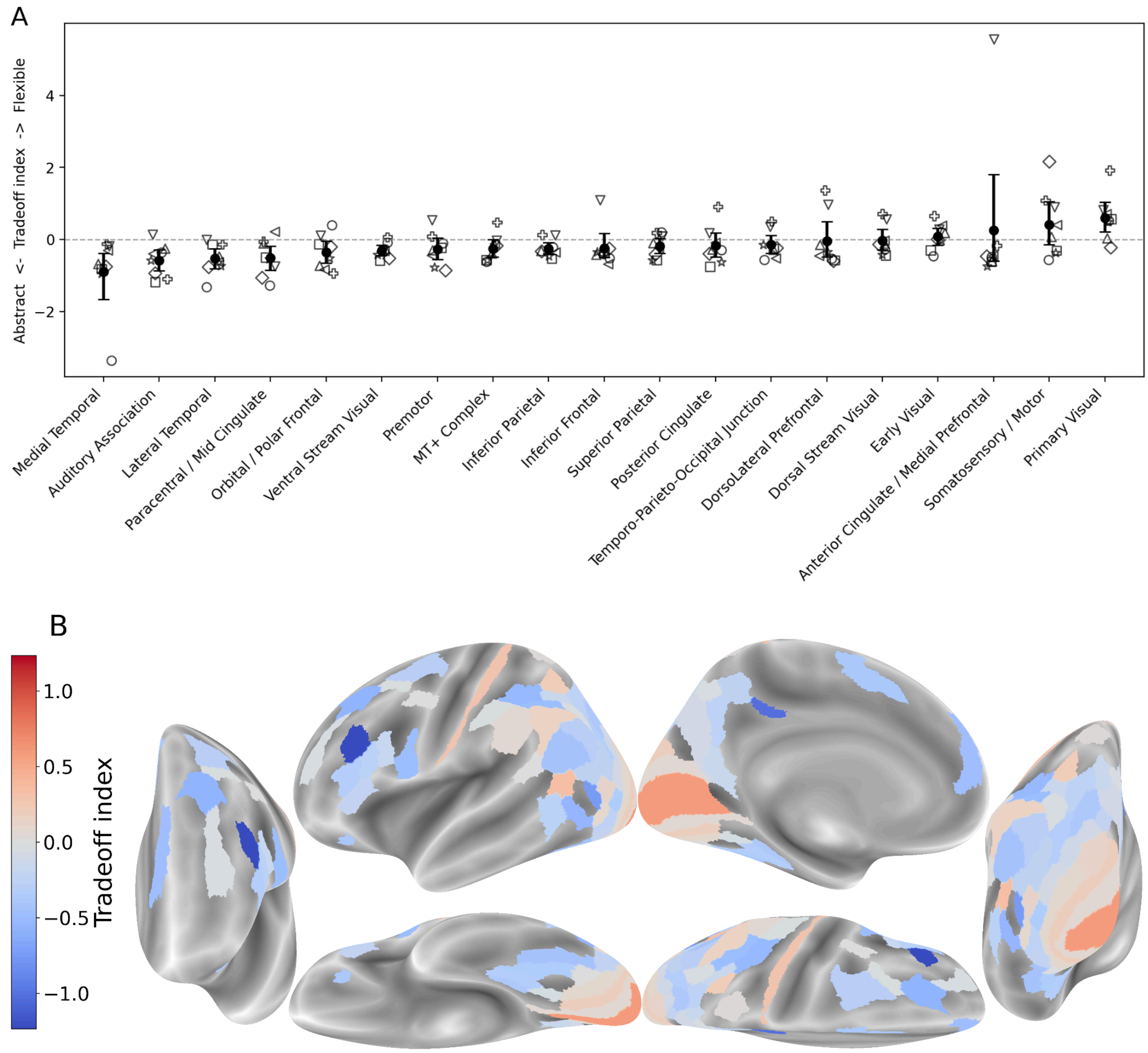


***Figure S11.Tradeoff index based on the SNR-independent measure (Equation S2).*** *The tradeoff index (A) for each of the 22 coarse-grained ROIs, and (B) for the fine-grained cortical atlas. The conventions are the same as in Figure 4. Overall, even when we use a more SNR-independent index of the tradeoff, the general trend remains similar: early sensory regions favored format flexibility at the cost of representational stability, whereas association cortex (particularly temporal regions) prioritized stability.*

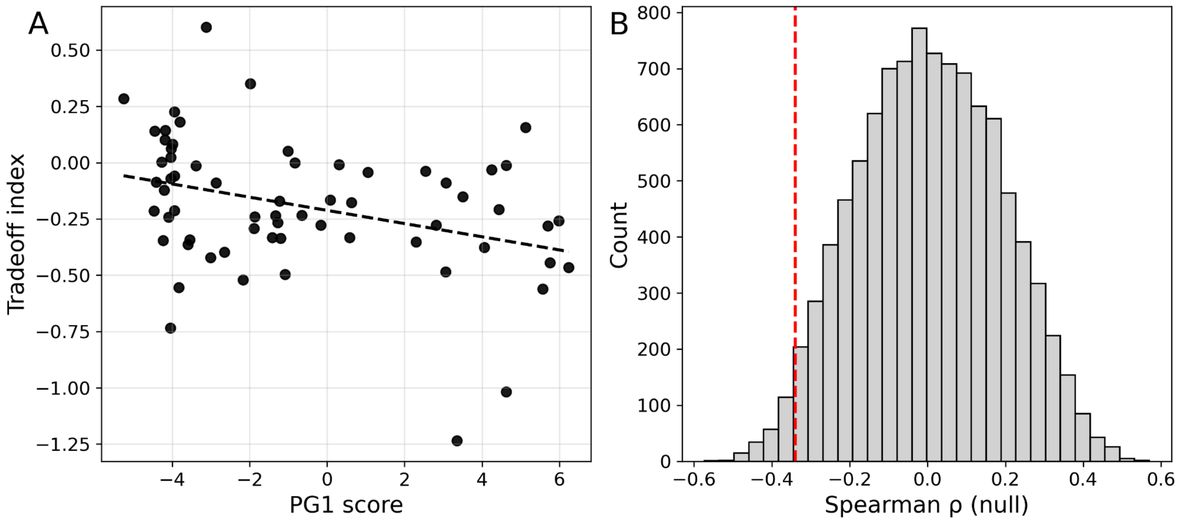


***Figure S12. Relationship between the tradeoff index (based on the SNR-independent measure) and the functional gradient.*** *(A) Tradeoff index plotted against the 1st principal gradient scores from Margulies et al. [s2] for the ROIs selected using the same criteria as in Figure 4. The Spearman correlation was significantly negative when tested against a null hypothesis assuming fully random spatial structure ($\rho = -.339$, two-sided $p = .006$). (B) Null distribution of Spearman ρ obtained while preserving spatial autocorrelation [s3, s4]. Under this more stringent null model, the observed correlation was barely non-significant (two-sided $p = .056$). The red vertical line indicates the observed correlation coefficient.*

### Supplementary tables

#### *Table S1. Group-level statistics for the rotation angle analysis.*

| ROI | Mean z | 95% CI | Cohen's dz | t | df | p | | q |
|---|---|---|---|---|---|---|---|---|
| Primary Visual | 2.76 | 1.58–4.07 | 1.39 | 3.93 | 7 | 0.003 | ** | 0.006 |
| Early Visual | 4.15 | 2.68–5.68 | 1.76 | 4.98 | 7 | $8.0\times10^{-4}$ | ** | 0.004 |
| Dorsal Stream Visual | 2.55 | 1.27–4.00 | 1.21 | 3.42 | 7 | 0.006 | * | 0.010 |
| Ventral Stream Visual | 3.21 | 2.16–4.39 | 1.84 | 5.20 | 7 | $6.3\times10^{-4}$ | ** | 0.003 |
| MT+ Complex | 2.57 | 1.57–3.61 | 1.64 | 4.64 | 7 | 0.001 | ** | 0.004 |
| Somatosensory / Motor | 0.54 | -0.18–1.33 | 0.46 | 1.31 | 7 | 0.116 | | 0.122 |
| Paracentral / Mid Cingulate | 0.90 | 0.02–1.75 | 0.67 | 1.90 | 7 | 0.049 | | 0.054 |
| Premotor | 1.25 | 0.35–2.06 | 0.94 | 2.67 | 7 | 0.016 | * | 0.024 |
| Posterior Opercular | 0.07 | -0.44–0.63 | 0.09 | 0.25 | 7 | 0.404 | | 0.404 |
| Early Auditory | 0.73 | 0.26–1.16 | 1.05 | 2.97 | 7 | 0.010 | * | 0.016 |
| Auditory Association | 0.49 | 0.11–0.89 | 0.81 | 2.28 | 7 | 0.028 | * | 0.037 |
| Insular / Frontal Opercular | 0.89 | 0.21–1.66 | 0.79 | 2.23 | 7 | 0.031 | * | 0.037 |
| Medial Temporal | 1.80 | 1.07–2.61 | 1.52 | 4.29 | 7 | 0.002 | ** | 0.005 |
| Lateral Temporal | 0.98 | 0.09–1.81 | 0.73 | 2.07 | 7 | 0.039 | * | 0.045 |
| Temporo-Parieto-Occipital Junction | 1.42 | 0.28–2.56 | 0.80 | 2.25 | 7 | 0.030 | * | 0.037 |
| Superior Parietal | 1.59 | 0.81–2.41 | 1.27 | 3.60 | 7 | 0.004 | ** | 0.009 |
| Inferior Parietal | 1.59 | 0.87–2.29 | 1.44 | 4.08 | 7 | 0.002 | ** | 0.006 |
| Posterior Cingulate | 2.48 | 1.70–3.28 | 1.99 | 5.63 | 7 | $4.0\times10^{-4}$ | ** | 0.003 |
| Anterior Cingulate / Medial Prefrontal | 1.42 | 1.06–1.81 | 2.43 | 6.86 | 7 | $1.2\times10^{-4}$ | ** | 0.003 |
| Orbital / Polar Frontal | 1.20 | 0.72–1.68 | 1.61 | 4.55 | 7 | 0.001 | ** | 0.004 |
| Inferior Frontal | 1.45 | 0.63–2.30 | 1.12 | 3.17 | 7 | 0.008 | * | 0.013 |
| DorsoLateral Prefrontal | 1.58 | 1.03–2.15 | 1.85 | 5.23 | 7 | $6.1\times10^{-4}$ | ** | 0.003 |

*Interobserver mean z-values (n = 8) are shown with 95% CIs and effect sizes (Cohen's dz = std / mean). t-values are shown with degrees of freedom (df), uncorrected p-values, and FDR-corrected q-values (** q < .01, * q < .05).*

#### *Table S2. Group-level statistics for the decoding analysis (Face vs Scene).*

| ROI | Mean z | 95% CI | Cohen's dz | t | df | p | | q |
|---|---|---|---|---|---|---|---|---|
| Primary Visual | 7.03 | 4.98–8.83 | 2.39 | 6.76 | 7 | $1.3\times10^{-4}$ | *** | $4.1\times10^{-4}$ |
| Early Visual | 12.81 | 9.97–15.64 | 2.85 | 8.07 | 7 | $< 1\times10^{-4}$ | *** | $3.5\times10^{-4}$ |
| Dorsal Stream Visual | 11.43 | 8.53–14.37 | 2.54 | 7.18 | 7 | $< 1\times10^{-4}$ | *** | $4.0\times10^{-4}$ |
| Ventral Stream Visual | 12.27 | 9.13–15.40 | 2.58 | 7.31 | 7 | $< 1\times10^{-4}$ | *** | $4.0\times10^{-4}$ |
| MT+ Complex | 12.38 | 9.61–15.30 | 2.81 | 7.95 | 7 | $< 1\times10^{-4}$ | *** | $3.5\times10^{-4}$ |
| Somatosensory / Motor | 2.97 | 1.45–4.58 | 1.23 | 3.47 | 7 | 0.005 | ** | 0.006 |
| Paracentral / Mid Cingulate | 3.03 | 1.33–4.54 | 1.22 | 3.45 | 7 | 0.005 | ** | 0.006 |
| Premotor | 8.04 | 5.31–10.87 | 1.86 | 5.27 | 7 | $5.8\times10^{-4}$ | *** | $9.1\times10^{-4}$ |
| Posterior Opercular | 1.97 | 1.28–2.61 | 1.95 | 5.53 | 7 | $4.4\times10^{-4}$ | *** | $8.8\times10^{-4}$ |
| Early Auditory | 0.53 | 0.03–1.27 | 0.54 | 1.52 | 7 | 0.087 | | 0.087 |
| Auditory Association | 3.36 | 2.04–4.95 | 1.48 | 4.18 | 7 | 0.002 | ** | 0.003 |
| Insular / Frontal Opercular | 2.28 | 0.97–3.59 | 1.12 | 3.16 | 7 | 0.008 | ** | 0.008 |
| Medial Temporal | 6.79 | 4.47–9.10 | 1.90 | 5.37 | 7 | $5.2\times10^{-4}$ | *** | $8.8\times10^{-4}$ |
| Lateral Temporal | 7.12 | 4.65–9.91 | 1.74 | 4.93 | 7 | $8.5\times10^{-4}$ | ** | 0.001 |
| Temporo-Parieto-Occipital Junction | 6.86 | 5.13–8.74 | 2.46 | 6.95 | 7 | $1.1\times10^{-4}$ | *** | $4.0\times10^{-4}$ |
| Superior Parietal | 11.37 | 7.76–14.91 | 2.04 | 5.78 | 7 | $3.4\times10^{-4}$ | *** | $7.5\times10^{-4}$ |
| Inferior Parietal | 11.15 | 8.72–13.42 | 3.05 | 8.62 | 7 | $< 1\times10^{-4}$ | *** | $3.5\times10^{-4}$ |
| Posterior Cingulate | 9.02 | 6.48–11.54 | 2.28 | 6.45 | 7 | $1.7\times10^{-4}$ | *** | $4.8\times10^{-4}$ |
| Anterior Cingulate / Medial Prefrontal | 4.89 | 3.14–6.82 | 1.73 | 4.88 | 7 | $8.9\times10^{-4}$ | ** | 0.001 |
| Orbital / Polar Frontal | 4.09 | 2.56–5.63 | 1.71 | 4.84 | 7 | $9.3\times10^{-4}$ | ** | 0.001 |
| Inferior Frontal | 7.68 | 5.25–10.36 | 1.93 | 5.45 | 7 | $4.8\times10^{-4}$ | *** | $8.8\times10^{-4}$ |
| DorsoLateral Prefrontal | 7.27 | 5.10–9.43 | 2.15 | 6.07 | 7 | $2.5\times10^{-4}$ | *** | $6.2\times10^{-4}$ |

*Interobserver mean z-values (n = 8) are shown with 95% CIs and effect sizes (Cohen's dz = std / mean). t-values are shown with degrees of freedom (df), uncorrected p-values, and FDR-corrected q-values (*** q < .001, ** q < .01).*

*Table S3. Group-level statistics for the decoding analysis (CCGP).*

| ROI | Mean z | 95% CI | Cohen's dz | t | df | p | | q |
|---|---|---|---|---|---|---|---|---|
| Primary Visual | 1.72 | 0.09–3.63 | 0.63 | 1.78 | 7 | 0.059 | | 0.065 |
| Early Visual | 7.19 | 5.11–9.37 | 2.16 | 6.10 | 7 | $2.5\times10^{-4}$ | ** | 0.001 |
| Dorsal Stream Visual | 7.10 | 4.35–9.89 | 1.63 | 4.62 | 7 | 0.001 | ** | 0.003 |
| Ventral Stream Visual | 9.23 | 6.52–11.97 | 2.20 | 6.21 | 7 | $2.2\times10^{-4}$ | ** | 0.001 |
| MT+ Complex | 9.23 | 6.70–11.85 | 2.31 | 6.53 | 7 | $1.6\times10^{-4}$ | ** | 0.001 |
| Somatosensory / Motor | 0.81 | -0.12–1.87 | 0.53 | 1.49 | 7 | 0.090 | | 0.094 |
| Paracentral / Mid Cingulate | 2.07 | 1.16–2.99 | 1.43 | 4.03 | 7 | 0.002 | ** | 0.005 |
| Premotor | 5.96 | 3.50–8.44 | 1.55 | 4.38 | 7 | 0.002 | ** | 0.004 |
| Posterior Opercular | 1.22 | 0.51–2.07 | 1.01 | 2.85 | 7 | 0.012 | * | 0.015 |
| Early Auditory | 0.28 | -0.42–0.83 | 0.29 | 0.82 | 7 | 0.220 | | 0.220 |
| Auditory Association | 2.15 | 0.98–3.37 | 1.17 | 3.31 | 7 | 0.006 | ** | 0.008 |
| Insular / Frontal Opercular | 0.93 | 0.08–1.73 | 0.73 | 2.07 | 7 | 0.039 | * | 0.045 |
| Medial Temporal | 5.59 | 4.06–7.04 | 2.43 | 6.86 | 7 | $1.2\times10^{-4}$ | ** | 0.001 |
| Lateral Temporal | 5.16 | 3.09–7.69 | 1.45 | 4.10 | 7 | 0.002 | ** | 0.005 |
| Temporo-Parieto-Occipital Junction | 4.61 | 3.08–6.34 | 1.81 | 5.12 | 7 | $6.8\times10^{-4}$ | ** | 0.003 |
| Superior Parietal | 8.16 | 5.11–11.48 | 1.67 | 4.71 | 7 | 0.001 | ** | 0.003 |
| Inferior Parietal | 7.81 | 5.59–10.10 | 2.20 | 6.22 | 7 | $2.2\times10^{-4}$ | ** | 0.001 |
| Posterior Cingulate | 6.04 | 3.40–8.55 | 1.50 | 4.25 | 7 | 0.002 | ** | 0.004 |
| Anterior Cingulate / Medial Prefrontal | 3.07 | 1.45–4.74 | 1.18 | 3.32 | 7 | 0.006 | ** | 0.008 |
| Orbital / Polar Frontal | 2.87 | 1.42–4.48 | 1.20 | 3.39 | 7 | 0.006 | ** | 0.008 |
| Inferior Frontal | 5.43 | 3.09–8.07 | 1.40 | 3.96 | 7 | 0.003 | ** | 0.005 |
| DorsoLateral Prefrontal | 4.52 | 2.18–6.59 | 1.32 | 3.74 | 7 | 0.004 | ** | 0.006 |

*Interobserver mean z-values (n = 8) are shown with 95% CIs and effect sizes (Cohen's dz = std / mean). t-values are shown with degrees of freedom (df), uncorrected p-values, and FDR-corrected q-values (** q < .01, * q < .05).*

#### *Table S4. Group-level statistics for the decoding analysis (XOR).*

| ROI | Mean z | 95% CI | Cohen's dz | t | df | p | | q |
|---|---|---|---|---|---|---|---|---|
| Primary Visual | 5.20 | 3.65–6.78 | 2.11 | 5.98 | 7 | $2.8\times10^{-4}$ | *** | $8.7\times10^{-4}$ |
| Early Visual | 8.47 | 5.80–10.99 | 2.12 | 5.98 | 7 | $2.8\times10^{-4}$ | *** | $8.7\times10^{-4}$ |
| Dorsal Stream Visual | 5.94 | 4.07–7.58 | 2.24 | 6.34 | 7 | $2.0\times10^{-4}$ | *** | $8.7\times10^{-4}$ |
| Ventral Stream Visual | 4.98 | 3.37–6.69 | 1.91 | 5.40 | 7 | $5.0\times10^{-4}$ | ** | 0.001 |
| MT+ Complex | 5.58 | 3.88–7.43 | 2.03 | 5.75 | 7 | $3.5\times10^{-4}$ | *** | $9.6\times10^{-4}$ |
| Somatosensory / Motor | 1.50 | 1.12–1.99 | 2.23 | 6.30 | 7 | $2.0\times10^{-4}$ | *** | $8.7\times10^{-4}$ |
| Paracentral / Mid Cingulate | 1.19 | 0.42–2.07 | 0.95 | 2.68 | 7 | 0.016 | * | 0.018 |
| Premotor | 3.16 | 1.85–4.40 | 1.59 | 4.50 | 7 | 0.001 | ** | 0.003 |
| Posterior Opercular | 0.60 | 0.25–0.98 | 1.07 | 3.02 | 7 | 0.010 | * | 0.013 |
| Early Auditory | 0.57 | 0.18–0.96 | 0.95 | 2.68 | 7 | 0.016 | * | 0.018 |
| Auditory Association | 0.53 | -0.42–1.57 | 0.35 | 0.98 | 7 | 0.181 | | 0.181 |
| Insular / Frontal Opercular | 0.58 | 0.19–1.04 | 0.88 | 2.49 | 7 | 0.021 | * | 0.022 |
| Medial Temporal | 1.61 | 0.83–2.51 | 1.24 | 3.51 | 7 | 0.005 | ** | 0.008 |
| Lateral Temporal | 1.57 | 0.49–2.66 | 0.91 | 2.57 | 7 | 0.018 | * | 0.020 |
| Temporo-Parieto-Occipital Junction | 3.08 | 2.49–3.79 | 3.10 | 8.77 | 7 | $< 1\times10^{-4}$ | *** | $5.6\times10^{-4}$ |
| Superior Parietal | 4.85 | 3.57–6.14 | 2.47 | 6.97 | 7 | $1.1\times10^{-4}$ | *** | $7.9\times10^{-4}$ |
| Inferior Parietal | 4.41 | 3.39–5.45 | 2.77 | 7.82 | 7 | $< 1\times10^{-4}$ | *** | $5.8\times10^{-4}$ |
| Posterior Cingulate | 3.61 | 2.43–4.88 | 1.93 | 5.47 | 7 | $4.7\times10^{-4}$ | ** | 0.001 |
| Anterior Cingulate / Medial Prefrontal | 1.02 | 0.39–1.73 | 0.97 | 2.75 | 7 | 0.014 | * | 0.018 |
| Orbital / Polar Frontal | 0.83 | 0.30–1.29 | 1.09 | 3.08 | 7 | 0.009 | * | 0.013 |
| Inferior Frontal | 2.95 | 1.78–4.29 | 1.53 | 4.32 | 7 | 0.002 | ** | 0.003 |
| DorsoLateral Prefrontal | 2.65 | 1.83–3.67 | 1.86 | 5.25 | 7 | $5.9\times10^{-4}$ | ** | 0.001 |

*Interobserver mean z-values ($n = 8$) are shown with 95% CIs and effect sizes (Cohen’s dz = std / mean). t-values are shown with degrees of freedom (df), uncorrected p-values, and FDR-corrected q-values (*** $q < .001$, ** $q < .01$, * $q < .05$).*
